# Loss of adaptive capacity in asthmatics revealed by biomarker fluctuation dynamics upon experimental rhinovirus challenge

**DOI:** 10.1101/684910

**Authors:** Anirban Sinha, René Lutter, Binbin Xu, Tamara Dekker, Barbara Dierdorp, Peter J. Sterk, Urs Frey, Edgar Delgado-Eckert

## Abstract

Asthma is a dynamic disease, in which lung mechanical and inflammatory processes often interact in a complex, unpredictable manner. We hypothesize that this may be explained by respiratory disease-related systems instability and loss of adaptability to changing environmental conditions, resulting in highly fluctuating biomarkers and symptoms. Using time series of inflammatory (eosinophils, neutrophils, FeNO), clinical and lung function biomarkers (PEF, FVC, and FEV_1_), we estimated this loss of adaptive capacity (AC) during an experimental perturbation with a rhinovirus in 24 healthy and asthmatic volunteers. Loss of AC was estimated by comparing similarities between pre- and post-challenge time series. Unlike healthy participants, the asthmatic’s post-viral-challenge state resembled significantly more other rhinovirus-infected asthmatics than their own pre-viral-challenge state (hypergeometric-test: p=0.029). This reveals loss of AC, and supports the novel concept that not only single physiological mechanisms, but interacting dynamic disease properties are altered in asthma and contribute to a more vulnerable phenotype.

## Introduction

The dynamics of (patho)physiology are particularly relevant in chronic diseases. In uncontrolled conditions, the patients are almost always clinically unstable, exhibiting a fluctuation pattern of symptoms, including unexpected loss of control or even sudden, potentially life-threatening exacerbations. Asthma, a chronic airway inflammation with variable airway obstruction is an archetypical example of such a dynamic disease. It is well known that the relationship between the strength of the environmental trigger, and the resulting degree of airway inflammation, airway obstruction, and symptoms is highly nonlinear, thus asthma is often insufficiently controlled with anti-inflammatory drugs (www.ginasthma.org) (1).

The temporal behavior of disease biomarkers in asthma is related to the inherently complex, interacting pathophysiology of the respiratory system, involving environmental, inflammatory and airway mechanical components (2). We and others have hypothesized that in health, the respiratory system exhibits a so-called ‘homeokinetic’ stable equilibrium state, enabling the respiratory system to easily adapt to an external perturbation (3). However, the homeokinesis of the respiratory system in asthma may be altered and adaptability to external stimuli (adaptive capacity) may be decreased, leading to a less stable system with more temporal fluctuations and exacerbations (3). Thus, asthma instability and fluctuation dynamics may not just relate to the degree of airway inflammation or the strength of the environmental trigger, but to the innate properties of the homeokinetic system itself.

Homeokinesis is the ability of a physiological system to return to a dynamic equilibrium after a perturbation (4–7); The remarkable complexity of such systems originates from non-linear interactions between their constitutive parts (8–10). The healthy homeokinetic respiratory system is characterized by a stable equilibrium and sufficient adaptability in the face of changing environmental conditions such as pollutants, pathogens and allergens (10). A concomitant phenomenon of homeokinesis is the fluctuation behavior of physiological processes and related physiological parameters and biomarkers around a stable medium value when measured longitudinally (11, 12). Over the past decades, a considerable research effort has been invested into mathematically analyzing and characterizing physiologic time series (8, 13–16). Moreover, it has been shown that time series properties, such as unpredictability, long-range correlations, fractality, and information content are associated with states of health and disease (10, 17). Such changes in dynamic behavior are potentially related to loss of adaptive capacity of the complex respiratory system. However, quantifying such loss of the system’s adaptive capacity, i.e., its ability to cope with external perturbations (10), is more difficult and dependent on the physiological context. For instance, researchers have directly linked the capacity of rats to adapt to environmental heat stresses to the ability of the animal’s liver cells to rapidly express the heat shock protein HSP70 in high quantities (18). Other scientists have suggested “the capacity of a physiological system to bring itself autonomously back to the normal homeostatic range after a challenge” as a more workable definition of adaptive capacity (19). We found evidence for a loss of adaptive capacity by quantitatively comparing the similarities of means and fluctuations in the pre- and post-viral-challenge time series of biomarkers in an unassuming data driven manner, using hierarchical clustering (20) and statistically assessing the clusters found by means of enrichment analysis (21–23).

The aim of this study is to test the hypothesis, whether in asthmatics the adaptive capacity to a standardized environmental perturbation, such as an experimental viral challenge, is altered in comparison to healthy subjects. As a proof of concept, such changes in the homeokinetic system properties in disease would support evidence that not only single factors, such as inflammation, but also system properties contribute to disease dynamics and phenotype stability.

In a prospective, longitudinally designed study comprising healthy and asthmatic subjects, we measured time series of a set of standard lung functional and inflammatory/immune biomarkers two months prior to and one month following an experimental rhinovirus 16 (RV16) infection induced by controlled and deliberate inoculation of healthy and asthmatic volunteers. This choice was driven by the fact that rhinovirus (RV) infections in asthmatics have been found to be among the most prominent external triggers of acute worsening of asthma symptoms, and of asthma exacerbations and loss of control (24, 25). Adaptive capacity to the rhinovirus infection was compared between asthmatic and healthy subjects for the abovementioned biomarkers.

## Results

### Experimental rhinovirus challenge while monitoring cohort participants

In all cohort participants (12 healthy and 12 asthmatic volunteers), the biomarkers/parameters listed in Table 1 below were measured during two months before, and during one month immediately after deliberate experimental inoculation with rhinovirus, resulting in pre- and post-viral-challenge time series of each biomarker/parameter. Plots of the time series of each biomarker can be found in Supplementary Figures File 2. For the healthy and the asthmatics groups separately, summary statistics of the average before the viral challenge (average over 2 months) and after the viral challenge (average over 1 month) of each of these biomarkers/parameters can be found in the SI.

**Table 1:** Biomarkers/parameters measured in each cohort participant during two months before, and during one month immediately after deliberate experimental inoculation with rhinovirus. The corresponding sampling frequencies can be found in columns 2 and 3. See the Methods section below for more details on the study design, and on the measurement procedures and laboratory assays used. FEV1: forced expiratory volume in one second. FVC: forced vital capacity. PEF: peak expiratory flow. FeNO: fractional expired concentration of nitric oxide.

| Biomarker or parameter | Sampling frequency before rhinovirus challenge | Sampling frequency after rhinovirus challenge |
| --- | --- | --- |
| Lung function (FEV1, FVC, FEV1/FVC, PEF) | 2x daily | 2x daily |
| Exhaled Nitric Oxide (FeNO) | 3x weekly | 3x weekly |
| Eosinophil and neutrophil cell density in nasal lavage fluid | 1x weekly | 3x weekly |

### Hierarchical clustering of biomarker time series

In order to quantitatively establish the degree of similarity or “proximity” between two time series of a given biomarker, we used the Earth Mover’s Distance (EMD), which regards each of the time series as a univariate empirical distribution of the biomarker at hand (see Methods and SI for more details). The pre- and post-challenge time series of individual biomarker time series from all participants (both healthy and asthmatics) were clustered using the EMD as the distance metric between the time series. The outcomes for the levels of exhaled nitric oxide (FeNO), and the percentage of eosinophils in nasal lavage fluid are presented here, whereas the results for the other biomarkers are presented in the SI.

#### i) Time series of exhaled Nitric Oxide (FeNO)

Findings are summarized in Table 2. The corresponding dendrogram is depicted in Figure 1. In brief, we found three clusters. Cluster 1 consists of four time series stemming from 2 asthmatics. As can be read off of the dendrogram in Fig. 1 below, and of the distance matrix depicted in Panel C of Fig. 2 (see Methods section below), these two participants are prominently different from the rest (regarding their FeNO time series), and might be regarded as outliers. Cluster 2 contains more healthy participants than expected by chance. In other words, Cluster 2 is *enriched* in healthy participants. Conversely, due to the balanced design of the cohort (equal numbers of healthy and of asthmatic participants), Cluster 2 is also *depleted* of asthmatic participants, i.e., it contains fewer asthmatic participants than expected by chance. And finally, Cluster 3, which is enriched in asthmatic participants. The marked difference between the members of Cluster 1 and Cluster 3 regarding FeNO time series revealed by the hierarchical clustering evidences a higher heterogeneity among the asthmatics, as compared to the healthy participants. In Cluster 2, the tendency for infected participants to be clustered together with their corresponding uninfected counterpart is statistically significant (p-value = 0.038, see Table 2 below). This is not the case for Cluster 3. The difference in this regard between Cluster 2 (mainly healthy participants) and Cluster 3 (mainly asthmatic participants) is further underpinned by the fact that the distributions of cophenetic distances between the infected cluster members and their uninfected counterparts are statistically significantly different between these two clusters (p-value=0.033, see also Fig. 3 in the SI). More specifically, an average of 34.5 in Cluster 3, as opposed to 12.0 in Cluster 2.

**Table 2:** Composition, enrichment analysis, and grouping characteristics of the clusters found by comparison of each participant’s pre- and post-challenge time series of FeNO. Enrichment is marked in bold letters, depletion in italics; the corresponding p-values were calculated using the hypergeometric test. The empirical p-values for the proportion of pre- and post-pairs were calculated using simulated permutations (see Methods section). A participant is fully represented in a given cluster if both their pre- and post-challenge time series of measurements are contained in the cluster. For example, the healthy participant “P08H” is fully represented in Cluster 2, as both their pre- and post-challenge time series of FeNO measurements are members of Cluster 2 (see Fig. 1 below). Partial representation corresponds to the scenario in which only one of the two time series (pre- and post-challenge) is a member of the cluster. For instance, the asthmatic participant “ P07A” is only partially represented in Cluster 2, because their pre-challenge time series of FeNO measurements is part of Cluster 2, whereas their post-challenge time series of FeNO belongs to Cluster 3 (see Fig. 1 below). See also the Methods section for the definition of neighbors.

| Characteristic \ Cluster number | Cluster 1 | Cluster 2 | Cluster 3 |
| --- | --- | --- | --- |
| **Size (%)** | 4 (8.33 %) | 26 (54.17 %) | 18 (37.5 %) |
| **Fully represented healthy participants** | 0 | 11 | 0 |
| **Partially represented healthy participants** | 0 | 1 | 1 |
| **Fully represented asthmatic participants** | 2 | 1 | 8 |
| **Partially represented asthmatic participants** | 0 | 1 | 1 |
| **Number of time series from healthy participants (%)** | 0 (0%) | 23 (88.46 %) | 1 (5.56 %) |
| **p-value** | 0.055 | **1.78E-09** | *1.15E-06* |
| **Number of neighboring pre- and post-pairs (%)** | 2 (100 %) | 4 (28.57 %) | 2 (20 %) |
| **Empirical p-value** | 0.003 | 0.007 | 0.097 |

**Figure 1:**
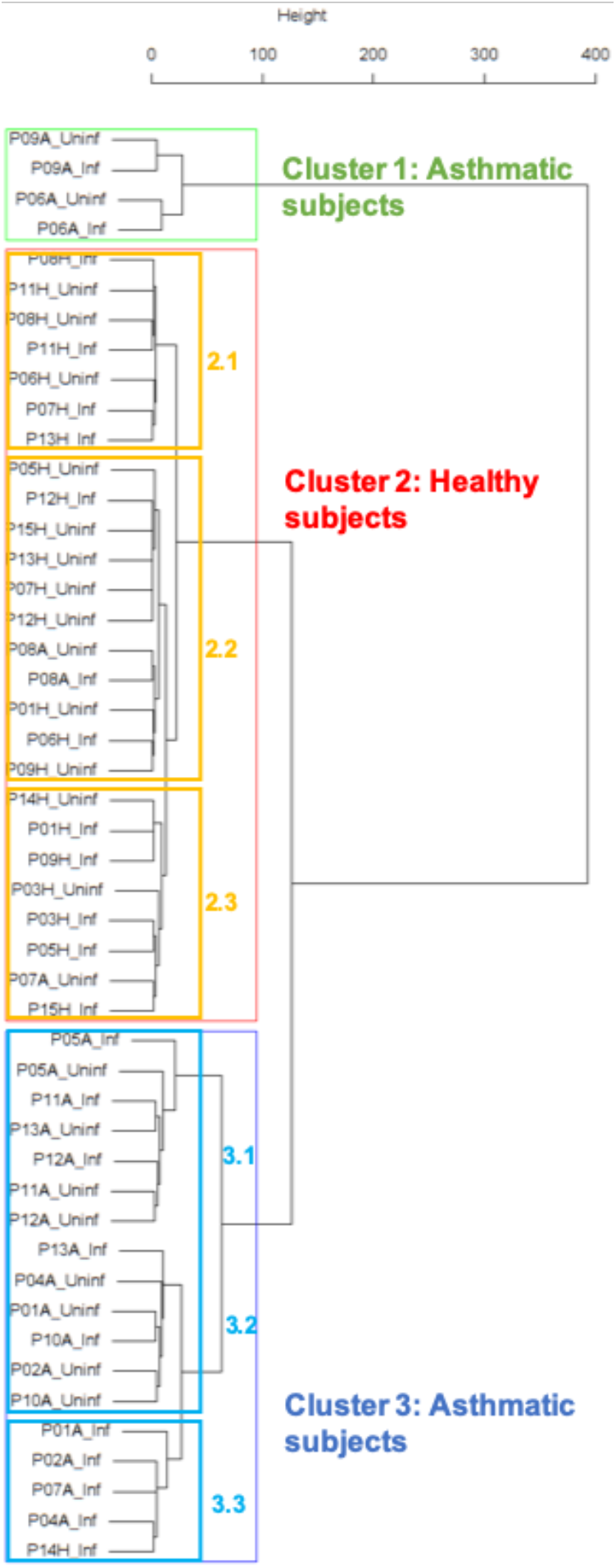
Cluster dendrogram obtained via hierarchical clustering of the participants’ pre- and post-challenge time series of FeNO. The distance between any two-time series was calculated using the EMD. Rectangles mark the clusters and sub-clusters identified. From top to bottom: Cluster 1, Cluster 2 (subdivided into Clusters 2.1, 2.2, and 2.3), and Cluster 3 (subdivided into Clusters 3.1 and 3.2, and 3.3). Patient IDs are indicated by Pxy, their health status using H/A, denoting Healthy or Asthmatic, and their RV infection status by Uninf/Inf, which stands for Uninfected/Infected. Cluster 1 consists of asthmatics which are prominently different from other asthmatic subjects in Cluster 3 and also from healthy subjects in Cluster 2. These might be regarded as outliers.

**Figure 2:**
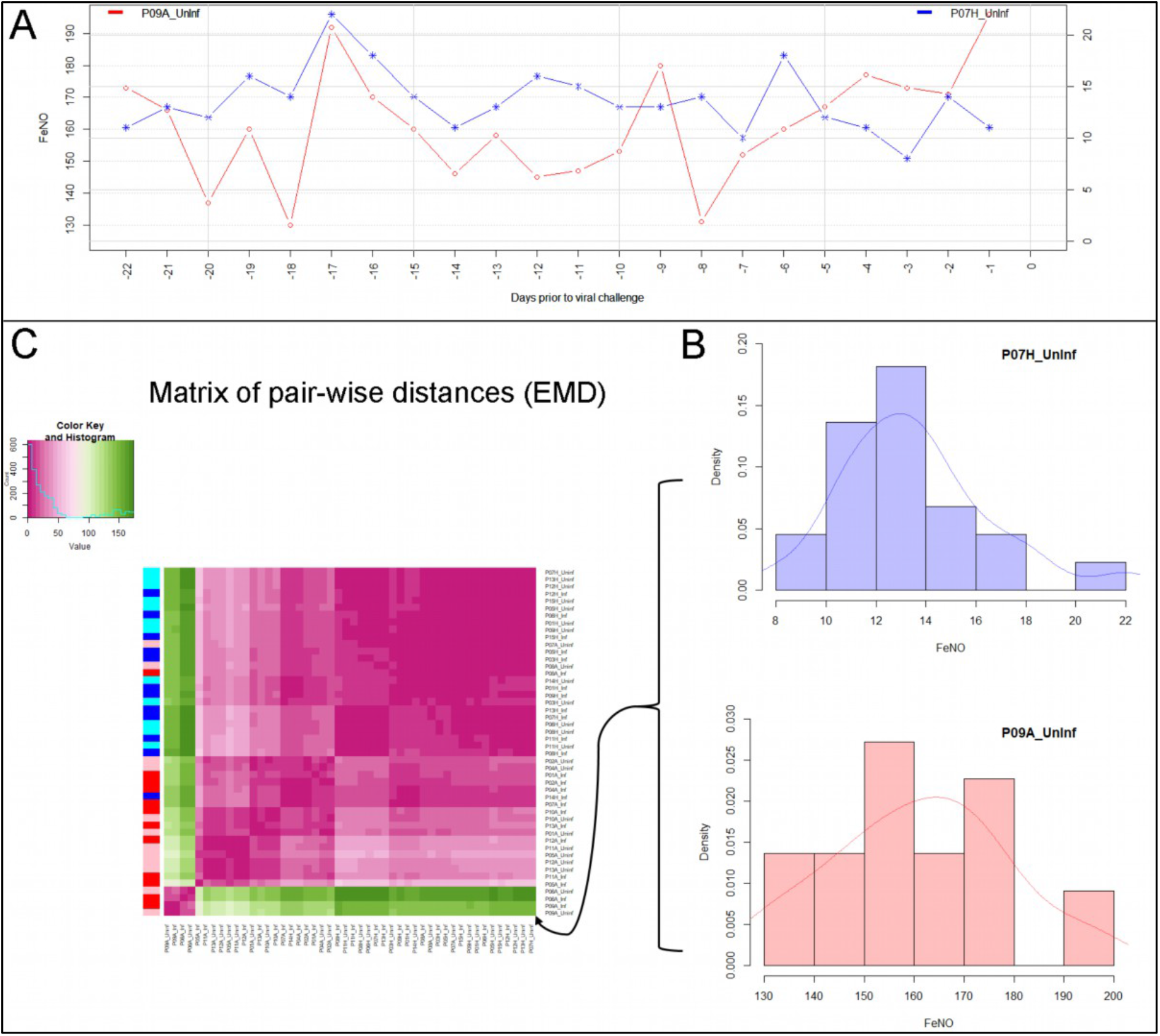
Panel A depicts two pre-challenge time series of FeNO obtained from a healthy (blue curve), and from an asthmatic (red curve) participant, respectively. In Panel B, each of the time series is represented as empirical distribution. This representation of the two-time series allows for the calculation of a distance or “dissimilarity” between the two by means of the Earth Mover’s Distance (EMD). The EMD-comparison of all possible pairs of time series (both pre- and post-challenge) results in a symmetric matrix of pair-wise distances, as shown in Panel C using a color-coded (violet to green) heat-map. Each row in this matrix corresponds to one time series. The color bar on the left hand side of the matrix encodes the “type” of time series: Cyan marks a pre-challenge time series originating from a healthy participant; Blue marks a post-challenge time series originating from a healthy participant; Pink marks a pre-challenge time series originating from an asthmatic participant; Red marks a post-challenge time series originating from an asthmatic participant. The information stored in the matrix of pair-wise distances is then used within an agglomerative clustering algorithm in order to group the time series in different clusters. The outcome of this procedure is represented using a dendrogram as depicted in Figure 1 above.

**Figure 3:**
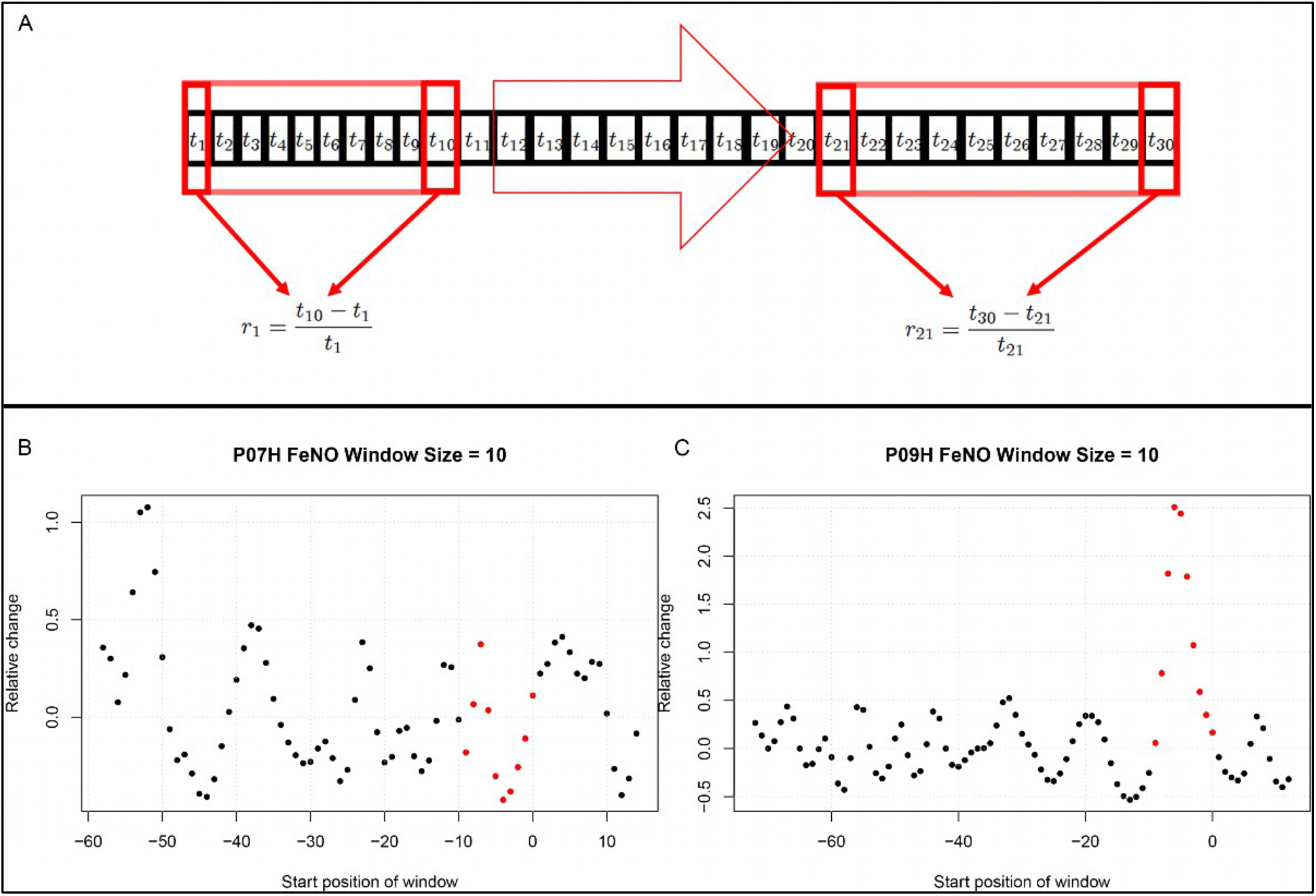
Panel A: Graphical representation of a biomarker time series *t*_*i*_. For the calculation of short-term/transient changes, a gliding interval or window is moved, one day at a time, along the time series. The relative change between the first and last entry of the gliding window is calculated, resulting in a new time series of short-term relative changes *r*_*i*_. In Panel B a healthy participant’s time series of short-term relative changes in FeNO is depicted. A gliding interval of size 10 days was used to calculate it from the participant’s time series of FeNO measurements. The start position of the gliding window is expressed relative to the day of the viral challenge, which is marked as day 0. When the position of the gliding window was such that the day of the viral challenge was contained within the gliding window, the corresponding value of the relative change is marked in red. In order to assess the statistical significance of the short-term relative changes possibly elicited by the viral challenge, the relative change values located to the left of those marked in red were compared to the values marked in red by means of a Mann-Whitney-U-test. Visual inspection of the time series in Panel B correctly suggests that the outcome of this test is not significant. The reason being that the relative changes within time intervals of 10 days observed prior to the viral challenge are comparable to changes observed within intervals of the same length containing the day of the viral challenge. In Panel C, depicting data from a different healthy participant, the situation is clearly different, as verified by a significant outcome of the corresponding Mann-Whitney-U-test. In such cases, the participant is called a “responder” with respect to the “relative change within 10 days criterion”.

In Table 3, the sub-clusters found within Clusters 2 and 3, respectively (marked with orange and blue rectangles in Figure 1), are analyzed in terms of enrichment in or depletion of pre- and post-challenge time series. This analysis provides evidence for a statistically significant separation of pre- and post-challenge time series within Cluster 3. Indeed, the union of subclusters 3.1 and 3.2 is enriched in pre-challenge time series (p-value=0.029, see Table 3 below), whereas subcluster 3.3 is enriched in post-challenge time series (p-value=0.029, see Table 3 below). Such a separation cannot be observed within Cluster 2.

**Table 3:** Enrichment analysis of the sub-clusters found within the clusters described in Table 2 above by comparison of each participant’s pre- and post-challenge time series of FeNO. Enrichment is marked in bold letters, depletion in italics; the corresponding p-values were calculated using the hypergeometric test.

| Sub-cluster number \ Characteristic | Size (%) | Number of pre-challenge series (%) | p-value |
| --- | --- | --- | --- |
| **Cluster 2.1** | 7 (26.92 %) | 3 (42.86 %) | 0.404 |
| **Cluster 2.2** | 11 (42.31 %) | 8 (72.73 %) | 0.104 |
| **Cluster 2.3** | 8 (30.77 %) | 3 (37.5 %) | 0.246 |
| **Cluster 3.1 & 3.2** | 13 (72.22 %) | 8 (61.54 %) | **0.029** |
| **Cluster 3.3** | 5 (27.78 %) | 0 (0.00 %) | *0.029* |

A bootstrap based sensitivity analysis of these findings can be found in the SI.

#### ii) Time series of percentage of eosinophils in nasal lavage fluid

Findings are summarized in Table 4. The corresponding dendrogram is depicted in Fig.1 in the SI. In brief, three clusters were identified. Cluster 1 consists of four time series stemming from 3 asthmatics; As can be read off of the dendrogram depicted in Fig. 1 in the SI, these time series are prominently different from all the other time series, and might be regarded as outliers. Cluster 2 is enriched in healthy participants. And finally, Cluster 3, which is enriched in asthmatic participants. As seen in the analysis of FeNO, the marked difference between the members of Cluster 1 and Cluster 3 evidences a higher heterogeneity among the asthmatics, as compared to the healthy participants. However, Cluster 1 in the eosinophil analysis and Cluster 1 in the FeNO analysis only have one asthmatic patient in common. Again, in Cluster 2, the tendency for infected participants to be clustered together with their corresponding uninfected counterpart is statistically significant (p-value = 0.001, see Table 4 below). This is not the case for clusters 1 and 3. The difference in this regard between Cluster 2 (mainly healthy participants) and Cluster 3 (mainly asthmatic participants) is further substantiated by the fact that the distributions of cophenetic distances between the infected cluster members and their uninfected counterparts are statistically significantly different between these two clusters (p-value= 8.96e-05, one-tailed Mann-Whitney-U-test, see also Fig. 2 in the SI). More specifically, an average of 36.0 in Cluster 3, as opposed to 0.5 in Cluster 2.

**Table 4:** Composition, enrichment analysis, and grouping characteristics of the clusters found by comparison of each participant’s pre- and post-challenge time series of percentage of eosinophils in nasal lavage fluid. Enrichment is marked in bold letters, depletion in italics; the corresponding p-values were calculated using the hypergeometric test. The empirical p-values for the proportion of pre- and post-pairs were calculated using simulated permutations (see Methods section). A participant is fully represented in a given cluster if both their pre- and post-challenge time series of measurements are contained in the cluster. Partial representation corresponds to the scenario in which only one of the two time series (pre- and post-challenge) is a member of the cluster. See also the Methods section for the definition of neighbors.

| Characteristic \ Cluster number | Cluster 1 | Cluster 2 | Cluster 3 |
| --- | --- | --- | --- |
| **Size (%)** | 4 (8.33 %) | 26 (54.17 %) | 18 (37.50 %) |
| **Fully represented healthy participants** | 0 | 11 | 1 |
| **Partially represented healthy participants** | 0 | 0 | 0 |
| **Fully represented asthmatic participants** | 1 | 2 | 7 |
| **Partially represented asthmatic participants** | 2 | 0 | 2 |
| **Number of time series from healthy participants (%)** | 0 (0%) | 22 (84.62 %) | 2 (11.11 %) |
| **p-value** | 0.055 | **1.09E-07** | 2.89E-05 |
| **Number of neighboring pre- and post-pairs (%)** | 1 (33.33 %) | 5 (38.46 %) | 1 (10 %) |
| **Empirical p-value** | 0.123 | 0.001 | 0.424 |

### Individual response to the viral challenge with respect to the biomarkers measured

In order to test the effectiveness of the virus challenge, we measured the individual patient’s response with respect to each predefined biomarker. The results of blood antibody tests (RV16 seroconversion) along with RV Polymerase Chain Reaction (PCR) conducted on nasal lavage fluid taken from every participant after the inoculation indicated that 11 out of 12 healthy participants and 12 out of 12 asthmatics were effectively infected with the RV16 after inoculation (Table 2 in the SI). According to these laboratory tests, one healthy participant did not become infected. However, this participant did develop cold symptoms within a few days after the virus inoculation, suggesting that the laboratory tests failed to detect the ongoing infection although the participant was positively infected. Consequently, this participant was included in the analyses.

After having established the efficacy of the inoculation with RV16, we then explored, for each of the biomarkers measured (listed in the first column of Table 5), for how many participants a statistically significant within-subject change upon infection can be observed (“responders”, see Table 5). To this end, two criteria for “responders” were implemented. The first criterion, which regards time series as univariate empirical distributions of the biomarker at hand, aimed at detecting distributional changes in a given biomarker induced by the viral challenge: Here, each participant’s pre- and post-challenge time series of each biomarker were compared using the Kolmogorov-Smirnov test. The second criterion aimed at detecting short-term and transient relative changes induced by the viral challenge in the context of the relative changes observed prior to the challenge. Here, throughout the entire period of observation, we assessed the relative change of each biomarker taking place within time intervals of 10 days. (see Subsection 5.2 and Fig. 3 in the Methods section below).

**Table 5:**
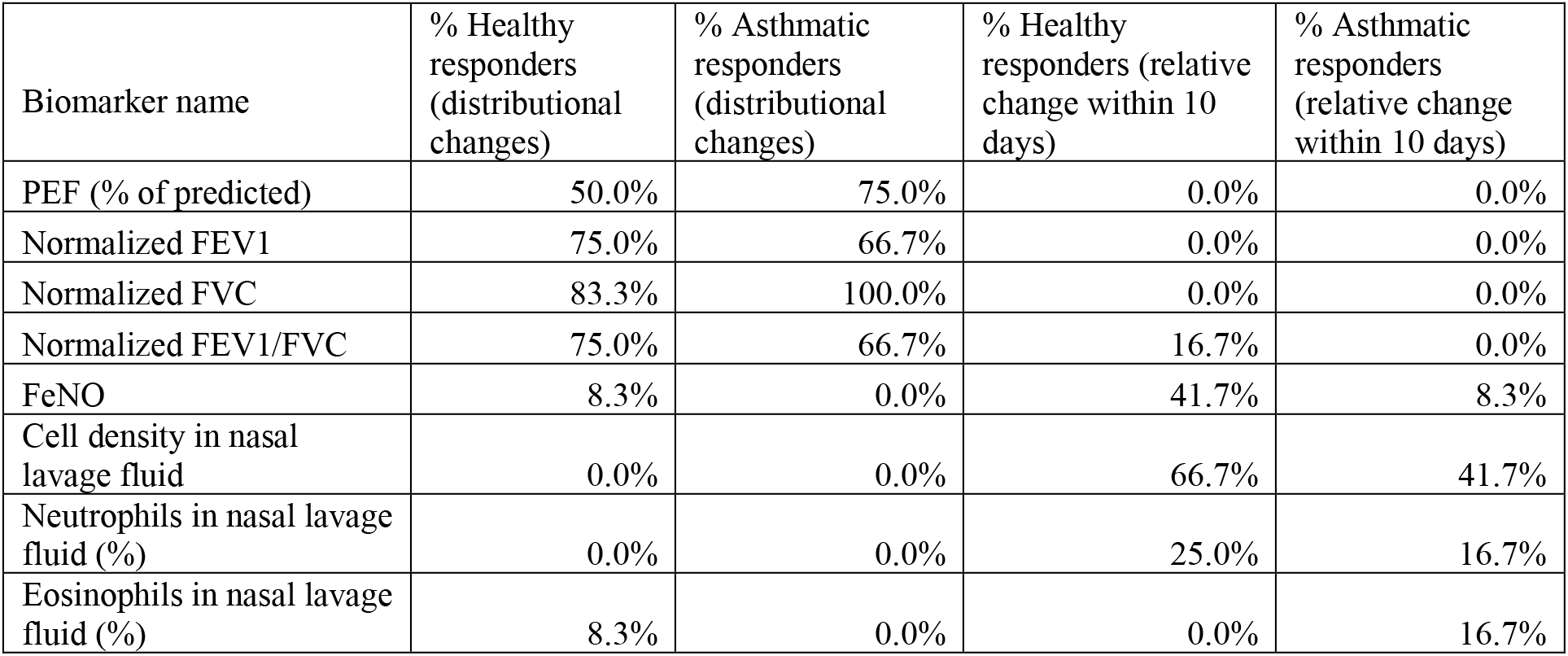
Proportions of responders within the groups of healthy and asthmatic participants, respectively. Two different criteria were used in order to establish a statistically significant response. According to the first criterion, a participant is considered a responder with respect to a given biomarker if the outcome of comparing the pre-challenge time series and the post-challenge time series of the same biomarker by means of the Kolmogorov-Smirnov test results in a p-value <= 0.05 (columns 2 and 3). According to the second criterion, a participant is considered a responder with respect to a given biomarker if the outcome of comparing, by means of a Mann-Whitney-U-test, the magnitude of relative changes observed during 10-day time intervals prior to the challenge with the magnitude of relative changes that took place during 10-day time intervals that contained the day of the challenge results in a p-value <= 0.05 (columns 4 and 5). For calculating the proportion of responders within each group the p-values were corrected for multiple testing using the false discovery rate (FDR) method of Benjamini and Hochberg. FEV1: forced expiratory volume in one second. FVC: forced vital capacity. PEF: peak expiratory flow. FeNO: fractional expired concentration of nitric oxide. The lung function parameters FEV1 and FVC, and thereby their ratio FEV1/FVC, were normalized using the standardized reference equations recommended by Global Lung Function Initiative (GLI) Task Force for comparisons across different populations.

## Discussion

In this proof of concept study, we provided experimental evidence for the loss of adaptive capacity in the human respiratory system due to asthma. To this end, we hypothesized that a loss of adaptive capacity could be experimentally demonstrated by means of a detection of a similarity diminution between the pre- and post-perturbation dynamics of the system. Using a data-driven clustering approach, we have shown that, in particular, FeNO and eosinophil time series were similar prior to and following the challenge in healthy subjects, suggesting stable homeokinetic behavior. In asthmatics, however, this similarity was predominantly reduced, suggesting a marked impact of the asthmatic condition on dynamic properties of the respiratory system, consistent with more unstable behavior and loss of adaptive capacity following the perturbation with viral infection. Adaptive capacity is a system characteristic of a homekinetic system. Our data on the respiratory system in asthma and health support the hypothesis that these homeokinetic system characteristics of the lung likely contribute to the stability and dynamic phenotype of asthmatic disease. This is particularly astonishing, since the average FeNO, lung function values, and nasal lavage fluid eosinophil counts are, in general, not different following a viral challenge.

### The notion of adaptive capacity in a clinical and epidemiological context

Epidemiological asthma research has provided indirect clues about a loss of adaptive capacity in the human respiratory system due to asthma. For instance, respiratory comorbidities have been found to be remarkably more prevalent in asthma than in non-asthma patients (26). Furthermore, provocation tests in asthma diagnostics (recently reviewed in (27)) implicitly rely on the assumption of loss of adaptive capacity in asthmatics. Loss of adaptive capacity manifests itself in increased vulnerability to external perturbations and e.g. the well-known fact that the common cold is a primary driver of asthma exacerbations (24, 28–38).

### Interpretation of our findings in a physiological context

Our study and findings for the first time demonstrate this at the system level using respiratory biomarker fluctuation dynamics. Loss of adaptive capacity cannot be determined by single-time point observations of biomarkers, it was, however, possible in a time series experiment involving asthma biomarkers during a standardized perturbation of the respiratory system. Nevertheless, the following considerations have to be discussed. We started off with the question: For which type of participant, healthy or asthmatic, and for which biomarker is the disruption introduced by the viral challenge strong enough to render infected individuals more similar among themselves than to their uninfected counterparts? Our results indicate that, with respect to cell density in nasal lavage fluid, there is a statistically significant separation of pre- and post-challenge time series. However, this effect does not seem to be cluster specific, even though we found a cluster of size 7 enriched in asthmatic participants within the subgroup of mainly post-challenge time series (see SI). A more distinctive scenario results from our cluster analysis of the pre- and post-challenge time series of the percentage of eosinophils in nasal lavage fluid: In Cluster 2, which is statistically significantly enriched in healthy participants, the tendency for infected participants to be clustered together with their corresponding uninfected counterpart is statistically significant and clearly higher than in Cluster 3, which is mainly composed of asthmatic participants. Finally, in the clustering of the pre- and post-challenge time series of FeNO we found Cluster 2, mainly composed of healthy participants, and Cluster 3, made of nearly 95% asthmatics. Furthermore, within Cluster 3 we found a statistically significant separation of pre- and post-challenge time series. No such separation was found within Cluster 2.

### Effectiveness of the perturbation (viral challenge)

Our analyses were systematically carried out with the general goal of testing the hypothesis of a loss of adaptive capacity of the respiratory system in asthmatic cohort participants. However, towards this goal, it was pertinent to determine whether and in which individuals the viral inoculation was effective. The presence of cold symptoms and/or positive blood antibody tests along with RV PCR provide strong evidence of all participants being effectively challenged shortly after RV16 inoculation. Indeed, RV PCR was performed on day 3 post inoculation and in all but one participant the presence of RV was detected. This is in concordance with other studies showing that RV concentrations in the nasal wash could be detected 2-3 days after viral inoculation, with a signal peak on the 5^th^/6^th^ day (39, 40).

### Variable and heterogeneous effect on lung function and inflammatory/immune biomarkers

We carried out a quantitative characterization of individual response to the viral perturbation. This was done using two computational/statistical approaches. One approach aimed to capture the changes elicited by the viral challenge taking place over longer time periods (comparison of the per- and post-challenge time series, viewed as empirical distributions), whereas the other assessed relative short-term changes occurring at shorter time scales (comparison of the magnitude of relative changes observed during 10-day time intervals).

There is a clear macroscopic/functional manifestation of the kindling RV infection, as reflected at the level of distributional changes induced by the viral challenge on lung function parameters. Indeed, with respect to this criterion, 50% or more statistically significant responders in each of the two groups (healthy and asthmatics) were found (see rows 1-4 in Table 5 above). Notably, significant differences found between pre- and post-challenge time series were, in general, not attributable to changes in the variance, as verified using Levene’s test (results not shown). Nevertheless, for most participants, *the lung function parameters* did not show short-term/transient relative changes induced by the viral challenge that were statistically significantly different in magnitude from short-term changes observed during the pre-challenge phase (see columns 4 and 5, and rows 1-4 in Table 5 above, and Figures in Supplementary Figures File 1). Taken together, these results suggest that the changes in lung function elicited by the viral challenge are, both for healthy and asthmatic participants, subtle, spread over comparatively longer time periods, and unlike a transient decline. This is in line with the results of previous studies (41–44), which concluded that after RV challenge lung function in asthmatic subjects did change, but did not decline dramatically in comparison to the changes observed in healthy controls. In contrast, our analyses indicate that changes in the inflammatory or immunological biomarkers at the cellular or molecular level are short-term and transient in nature (see rows 5-8 in Table 5 above, and Figures in Supplementary Figures File 1). Nevertheless, for these parameters fewer responders were found, when compared to the lung function parameters. However, our results also hint at a relatively short time scale of response of these inflammatory/immunological biomarkers. Thus, the sampling frequency used in this study may not entirely capture the rapidly changing magnitudes. The observed differences in the type of response between the lung function and the inflammatory/immunological biomarkers may be a manifestation of the interplay of different temporal and spatial scales.

### Limitations of the study

Our findings need to be judged in light of the limitations of our study. One of the limitations is that we only included mild asthmatics for ethical reasons. However, more severe asthmatics are likely to be on corticosteroid treatment, which may potentially influence the inflammatory response, therefore introducing a confounding factor. Another potential shortcoming of this study is the relatively small sample size. However, this drawback is compensated for by the unprecedented high sampling frequency at which the participants were screened in our study. In fact, the sample size was a compromise between the number of subjects and the number of assessment visits. Nevertheless, as discussed above, the sampling frequency for some of the inflammatory/immunological biomarkers may still not have been high enough in order to capture their temporal dynamics.

### Conclusion and implications

We have found evidence supporting the notion that a chronic disease such as asthma may alter the properties of a homeokinetic physiologic system in a way that compromises its capacity to appropriately react to a possibly harmful environmental stimulus. This loss of adaptive capacity in the asthmatic lung may be understood as changes that render the system overly unstable (3). As a proof of concept, such changes in homeokinetic system properties would support evidence that not only single factors, such as inflammation, but also system properties contribute to disease dynamics and phenotype stability.

This systems-level understanding of chronic asthma may open up new avenues for the understanding of asthma and other chronic dynamic diseases. Already in this small sample size, it is obvious that there is remarkable individual temporal variability in inflammatory and physiological biomarkers, not only in disease but also in health. Thus, dynamic fluctuations of physiological processes around an equilibrium state, including their related biomarkers, are an intrinsic feature of the respiratory system. Moreover, even in health there are strong inter-individual differences in these dynamic characteristics. Nevertheless, within a given healthy subject fluctuations remain similar following the virus challenge, indicating that, these dynamic fluctuations seem to be in a stable equilibrium state in health. This characteristic is lost in asthma.

Future studies involving time series of biomarker measurements may help us understand this system instability in chronic asthma, even in the absence of severe airway inflammation or obstruction. Furthermore, future therapeutic approaches may want to focus on maintaining a stable homeokinetic equilibrium of the respiratory system, rather than just normalizing single physiological or inflammatory biomarkers.

## Materials and Methods

This study was approved by the medical ethical committee from the Amsterdam University Medical Center and registered at the Netherlands Trial Register (NTR5426).

### 1. Participant cohort

Twelve non-smoking, atopic (as determined by positive skin prick test to common aero allergens), mild to moderate asthmatic subjects (based on ATS/ERS criteria), not using steroids were chosen for inclusion. Similarly, 12 non-smoking, non-atopic healthy subjects were also included in the study as controls. All participants provided written informed consent. The demographics of the study population are summarized in Table 7. All the participants were required to have their serum antibody titer of RV16 < 1:8 during screening. The age group for the study population was 18-30 years. Individuals with concomitant disease and pregnant women were excluded.

The basic inclusion criteria for the study populations followed standard recommendations and were as shown in Table 6 below.

**Table 6:**
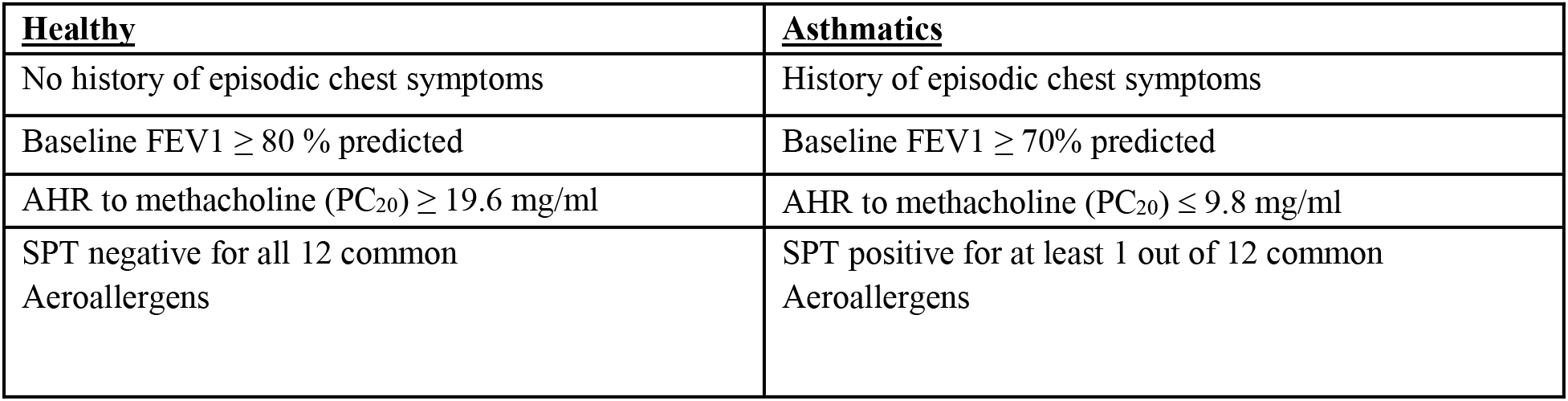
Basic characteristics of the study population. FEV1: forced expiratory volume in one second, AHR: Airway Hyper Responsiveness, PC_20_: Provocative Concentration causing a 20% fall in FEV1, SPT: Skin Prick Test

**Table 7:** The demographics of the study population. BMI is Body Mass Index. Only 1 healthy subject smoked 2 pack years or less 2 years before recruitment to our study, which is considered an insignificant smoking history. FEV1: forced expiratory volume in one second. PEF: peak expiratory flow.

| **Demographic features** | **Healthy** | **Asthmatic** |
| --- | --- | --- |
| Total number, n | 12 | 12 |
| Female gender, n (%) | 7 (58.3%) | 8 (66.7%) |
| Age (years), mean (SD) | 21 ± 1.5 | 22.2 ± 2.2 |
| Ethnicity (Caucasian), n (non-Caucasian, n) | 11 | 9 |
| BMI, mean (SD) | 22.2 ± 1.6 | 22.8 ± 3.1 |
| Smoking (pack years), n | 1 (0.17 PY) | -- |
| Height (centimeters) | 177.7 ± 8.6 | 172.5 ± 13.0 |
| Weight (KG) | 70.4 ± 10.1 | 67.8 ± 12.4 |
| Baseline spirometry |  |  |
| FEV1 %predicted | 105.7 ± 11.6 | 101.0 ± 10.0 |
| FVC %predicted | 104.2 ± 10.5 | 104.2 ± 10.2 |
| PEF %predicted | 108.4 ± 14.0 | 104.7 ± 12.2 |
| | | mean ± standard deviation |

### 2. Study Design

The project represents a prospective observational, follow-up study including patients with asthma and healthy controls with an experimental RV intervention.

The study participants were recruited after meticulous screening of volunteers (as mentioned in SI). The study was mainly divided into 2 phases. Phase 1 (stable phase) consisted of 2 months where these subjects were followed up and sampled diligently every alternate day (3 times a week for most of the measurements) with constant frequency at the hospital clinic. After that they were subjected to a standardized nasal dose of RV inoculation in the laboratory and followed up at the same frequency for one additional month (also called Phase 2 or unstable phase). In total the study consisted of 3 months of sampling period with a minimum of 180 measurements of lung function, 33 FeNO data points, and 20 cytokine and cell count measurements per subject.

The schematic work flow of the phases mentioned, is provided in the Fig 10 of the SI.

### 3. Measurement and collection of Biomarkers

#### 3.1 Lung function assessment

Spirometry was performed only once on the screening visit at the clinic to include participants based on inclusion criteria using a daily calibrated spirometer according to European Respiratory Society (ERS) recommendations (45).

Home monitoring of morning and evening lung function was done by hand held devices (Micro Diary, CareFusion, yielding the FEV1, FVC, FEV1/FVC and PEF values analyzed in this study. Moreover, the Asthma Control Questionnaire was administered.

#### 3.2 Exhaled Nitric Oxide (FeNO)

Measurement of fractionated exhaled nitric oxide (FENO) was performed using the NIOX MINO® (Aerocrine AB, Sweden). Single measurements per person were recorded at the clinic, thrice weekly, according to recommendations by the ATS (46).

#### 3.3 Nasal lavage

Nasal lavage was collected from the study participants once weekly before RV challenge and was up scaled to thrice weekly after the challenge at the clinic as previously described (42) [Refer to SI for details].

Table 8 provides an overview of the different sample measurements along with their frequency before and after rhinovirus challenge.

**Table 8:** The overview of different measurements performed in the study along with the frequency of sampling before and after rhino-virus challenge. Measures 1-4 include repeated measurements and 5,6 represent one-time measurement to screen the subjects for the study. 8 refers to the experimental intervention in the study. FEV1: forced expiratory volume in one second. FVC: forced vital capacity. PEF: peak expiratory flow. FeNO: fractional expired concentration of nitric oxide.

| Measurements of biomarkers | Frequency before rhinovirus challenge | Frequency after rhinovirus challenge |
| --- | --- | --- |
| Lung function with pocket-size spirometers (FEV1, FVC, FEV1/FVC, PEF) | 2x daily | 2x daily |
| Exhaled Nitric Oxide (FeNO) | 3x weekly | 3x weekly |
| Differential cell counts | 1x weekly | 3x weekly |
| Asthma Control Questionnaire | 2x daily | 2x daily |
| Spirometry | Performed once during screening to include subjects in the study |  |
| Methacholine challenge | Performed once during screening to include subjects in the study |  |
| Rhinovirus challenge | Performed after 2 months into the study |  |

### 4. Rhinovirus challenge

The study participants were exposed to rhinovirus 16 (RV16) using a standardized and validated challenge approach, based on previous studies by ourselves and other groups (47–52). All participants were screened for the presence of respiratory viruses just before the challenge, to rule out a concomitant infection resulting in a cold (see SI for more details). Those participants with a positive outcome of this test were excluded from the study. An experimental RV16 infection was induced by using a relatively low-dose inoculum of 100 TCID50 (Tissue Culture Infective Dose determining the amount of virus required to cause cytopathy in 50% of the cells) to mimic a natural exposure. The study protocol along with the viral dose used and its safety have been approved by the institutional Medical Ethics Committee in Amsterdam University Medical Centre, the details of which have been included in SI. Data from our previous study show that a low dose is sufficient to induce mild cold-symptoms (52, 53). Furthermore, this low-dose inoculum previously resulted in a slight decrease of FEV1 (loss of asthma control) in asthmatic patients between day 4 and 6 after RV16 exposure, whereas no decrease has been observed in healthy controls (50).

Refer to SI for further details.

### 5. Statistical and computational Analysis

Statistical tests resulting in a p-value less or equal to 0.05 were regarded as significant.

#### 5.1 Assessment of differences: Pre- vs. post-viral-challenge

For each participant, their time series of a given biomarker prior to and after the viral challenge were compared. This comparison was based on the Kolmogorov-Smirnov test, whereby the time series were treated as empirical distributions, thus disregarding the chronological order of the measurements.

Differences in the variance between the pre- and post-challenge distributions were assessed using Levene’s test (54).

Multiple comparison correction was performed where required, using the false discovery rate (FDR) method of Benjamini and Hochberg (55), setting the expected proportion of falsely rejected null hypotheses to 0.05.

The time series of a given biomarker, prior to and after the viral challenge, were regarded as empirical distributions and compared to each other using the Earth Mover’s Distance (EMD) (56). The resulting pair-wise distances between distributions were then used for hierarchical clustering of pre- and post-challenge distributions. See Fig. 2 below and the SI for more details. Our clustering approach makes use of the entire time series (distributions) of values measured before and after the challenge, respectively, and does not amalgamate the information into a single magnitude (e.g., the mean value). This method unveils subtle differences and similarities between the participants’ measurements that are less likely to be captured by conventional methods based on averages.

#### 5.2 Calculation of short-term/transient changes

For each participant individually, and for each biomarker, throughout the entire period of observation, the biomarker’s relative change in value taking place within time intervals of 10 days was calculated. This choice of time interval length was made based on published literature whereby 5 days post exposure to respiratory viruses was shown to be critical. Hence a 10-day window for comparison would include 5 days before challenge to contrast with 5 days after challenge (57). This was done throughout the entire period of observation considering all possible time intervals consisting of 10 consecutive days. In order to assess the statistical significance of the short-term relative changes possibly elicited by the viral challenge, the magnitude of relative changes observed during 10-day time intervals starting at least 10 days prior to the challenge were compared, by means of a Mann-Whitney-U-test, to the magnitude of relative changes that took place during 10-day time intervals that contained the day of the challenge. See Fig. 3 below and the SI for more details.

#### 5.3 Characterization of the dendrogram clusters

In order to evaluate the discriminatory power of a given biomarker, the clusters found in the clustering dendrogram were tested for enrichment in or depletion of healthy or asthmatic participants, and/or for enrichment in or depletion of pre- or post-challenge distributions. Statistically significant enrichment or depletion were established using the hypergeometric test (21–23).

The relative location of leaves in the clustering dendrogram was quantitatively evaluated using the cophenetic distance (58). The cophenetic distance between two leaves of a dendrogram is defined as the height of the dendrogram at which the two largest branches that individually contain the two leaves merge into a single branch.

For every cohort participant and any given biomarker there is a pre-challenge and a post-challenge time series, which we call the participant’s pre- and post-pair. If the disruption caused by the viral challenge is not strong enough, the pre- and post-challenge distributions of a given participant will tend to cluster together. Therefore, a cluster in which pre- and post-pairs are closely located in terms of the cophenetic distance within the dendrogram, represents a subgroup of participants for which the viral challenge caused a relatively weaker disruption, at least with respect to the biomarker under scrutiny.

Two dendrogram leaves are called neighbors if their mutual cophenetic distance is equal to the minimum of all cophenetic distances from one of the leaves to all the other leaves in the dendrogram. If this condition is fulfilled for both leaves simultaneously, then the two leaves form a two-element cluster in the dendrogram. If the condition is only fulfilled for one of the leaves, the two are still considered neighbors, even if this is not always visually obvious from inspecting the dendrogram (see Figure 11 in the SI).

Under the null-hypothesis that the branching in the dendrogram is the result of a purely random process, the number of neighboring pre- and post-pairs to be expected just by chance within a given cluster can be estimated by simply permuting the labels of the leaves in the dendrogram and counting the number of neighboring pre- and post-pairs. This permutation test is used for calculating the empirical p-values displayed in Tables 2 and 4 above.

A participant is fully represented in a given cluster if both their pre- and post-challenge time series of measurements are contained in the cluster. For example, the healthy participant “P08H” is fully represented in Cluster 2, as both their pre- and post-challenge time series of FeNO measurements are members of Cluster 2 (see Fig. 1 above). Partial representation corresponds to the scenario in which only one of the two time series (pre- and post-challenge) is a member of the cluster. For instance, the asthmatic participant “ P07A” is only partially represented in Cluster 2, because their pre-challenge time series of FeNO measurements is part of Cluster 2, whereas their post-challenge time series of FeNO belongs to Cluster 3 (see Fig. 1 above).

## Acknowledgments

The salary of AS was sponsored from the European Respiratory Society-Marie Sklodowska Curie actions COFUND RESPIRE 2 fellowships (MCF-7077-2014) and also from a grant supported by Lungenliga Schweiz (2017_14).

The work was supported by an unrestricted grant from Chiesi Pharmaceuticals, institutional funding from the Academic Medical Centre, Amsterdam UMC, University of Amsterdam (IA601011).

## Competing interests

None of the authors have any competing interest/s related to this manuscript.

## Supplementary Information

### Results

#### Effectiveness of the viral inoculation

Each participant in the study was administered the same dose of the virus (100 TCD 50) through the nose and every subject was tested for being positive for the virus after inoculation. False positive results due to previous exposure to the virus was ruled out by strict inclusion criteria of not having the titer of antibodies against RV16 > 1:8 in serum, measured at screening and prior to inoculation.

Positivity to viral inoculation was confirmed by either one of these three criteria

1. Positive test for antibodies against RV at the terminal visits of the participant
2. Positive RV PCR test from approximately 3^rd^ day (second visit) post RV challenge
3. Symptoms of RV induced cold

The effect of the virus in every study participant is summarized below in Table 1 (Column 1 indicates the seroconversion of the antibodies against the virus at the end of the study, Column 2 reflects the response to the virus by the PCR product on the day 3 after challenge and finally Column 3 shows the symptoms developed in the volunteers after the viral inoculation). Positive response in either of the 3 categories was considered evidence for a successful viral challenge.

**Table 1:**
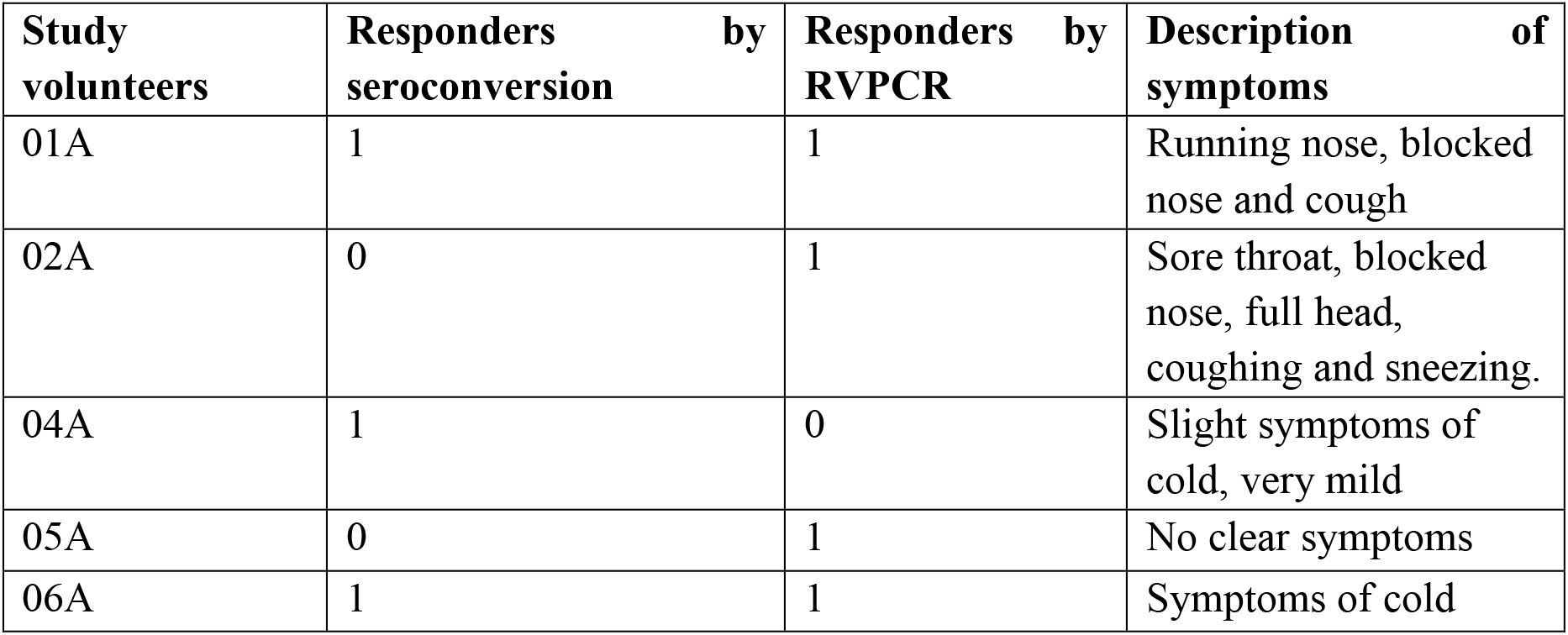
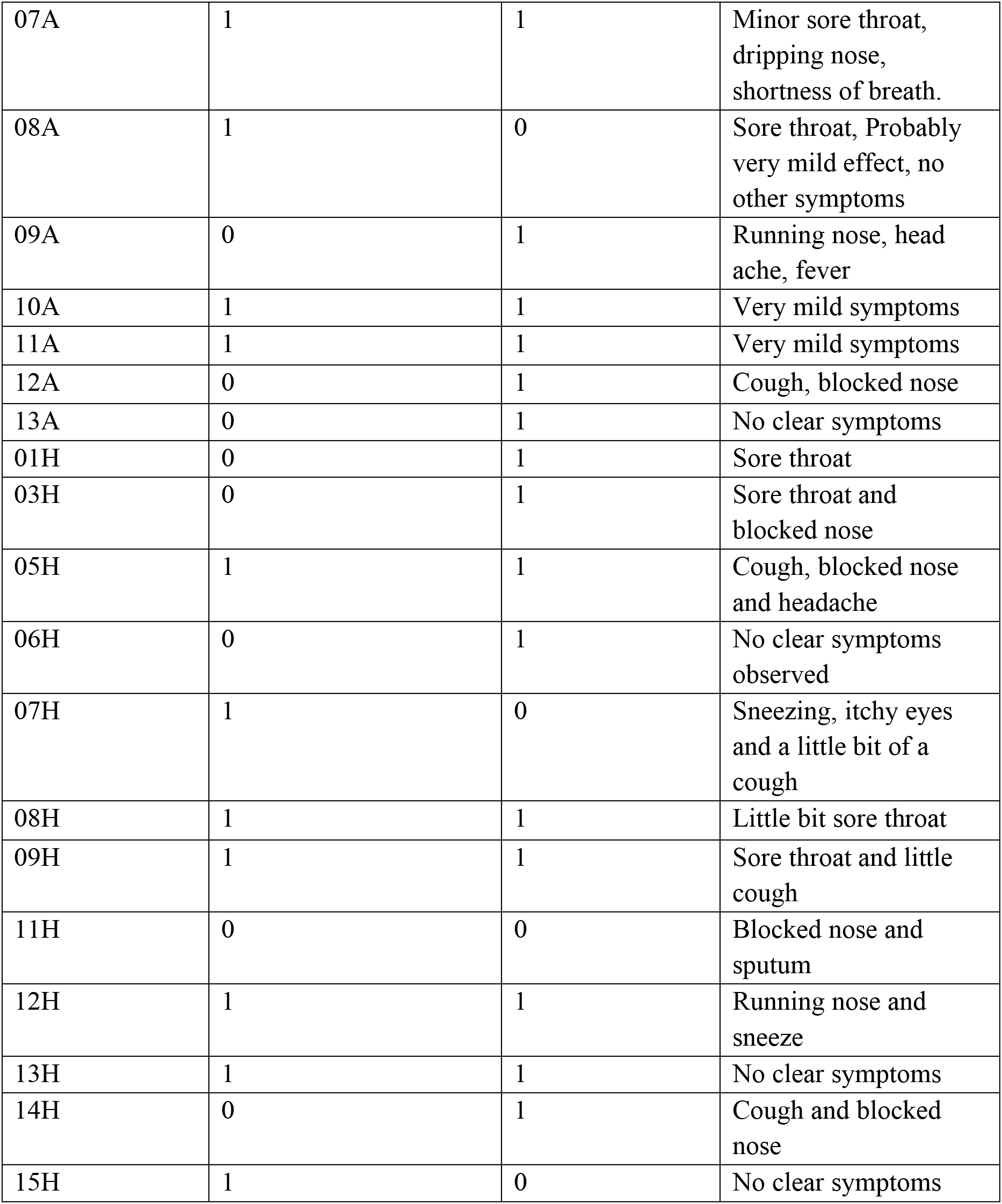
**Effectiveness of the viral challenge in** asthmatic and healthy participants, respectively. A participant is considered to have a successful viral inoculation if any one of the 3 tests is positive. 1 indicates positive response and 0 indicates a failed response in the corresponding tests indicated in columns.

| Study volunteers | Responders by seroconversion | Responders by RVPCR | Description of symptoms |
| --- | --- | --- | --- |
| 01A | 1 | 1 | Running nose, blocked nose and cough |
| 02A | 0 | 1 | Sore throat, blocked nose, full head, coughing and sneezing. |
| 04A | 1 | 0 | Slight symptoms of cold, very mild |
| 05A | 0 | 1 | No clear symptoms |
| 06A | 1 | 1 | Symptoms of cold |
| 07A | 1 | 1 | Minor sore throat, dripping nose, shortness of breath. |
| 08A | 1 | 0 | Sore throat, Probably very mild effect, no other symptoms |
| 09A | 0 | 1 | Running nose, head ache, fever |
| 10A | 1 | 1 | Very mild symptoms |
| 11A | 1 | 1 | Very mild symptoms |
| 12A | 0 | 1 | Cough, blocked nose |
| 13A | 0 | 1 | No clear symptoms |
| 01H | 0 | 1 | Sore throat |
| 03H | 0 | 1 | Sore throat and blocked nose |
| 05H | 1 | 1 | Cough, blocked nose and headache |
| 06H | 0 | 1 | No clear symptoms observed |
| 07H | 1 | 0 | Sneezing, itchy eyes and a little bit of a cough |
| 08H | 1 | 1 | Little bit sore throat |
| 09H | 1 | 1 | Sore throat and little cough |
| 11H | 0 | 0 | Blocked nose and sputum |
| 12H | 1 | 1 | Running nose and sneeze |
| 13H | 1 | 1 | No clear symptoms |
| 14H | 0 | 1 | Cough and blocked nose |
| 15H | 1 | 0 | No clear symptoms |

#### Clustering based on time series of biomarkers

##### Percentage of eosinophils in nasal lavage fluid

The corresponding dendrogram is depicted in Fig. 1 below.

**Figure 1:**
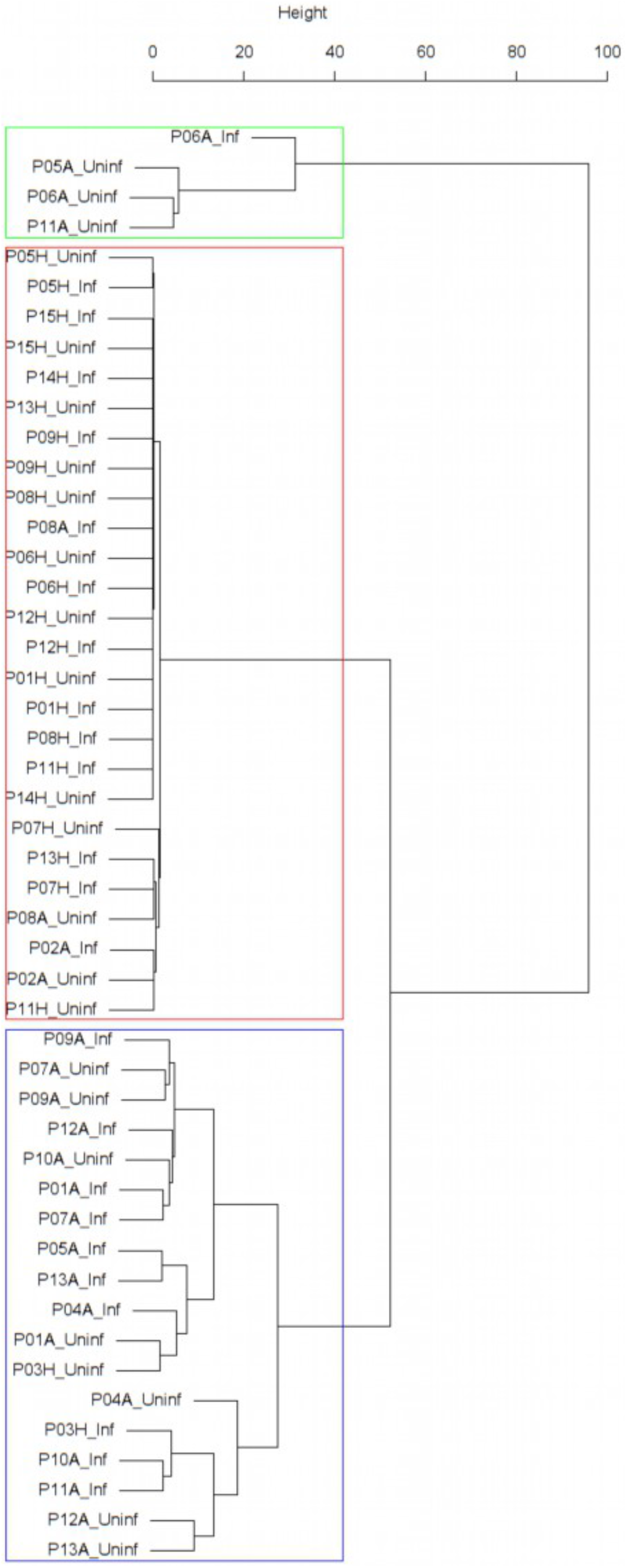
Dendrogram obtained from clustering the participants’ time series of the percentage of eosinophils in nasal lavage fluid using the EMD.

**Figure 2:**
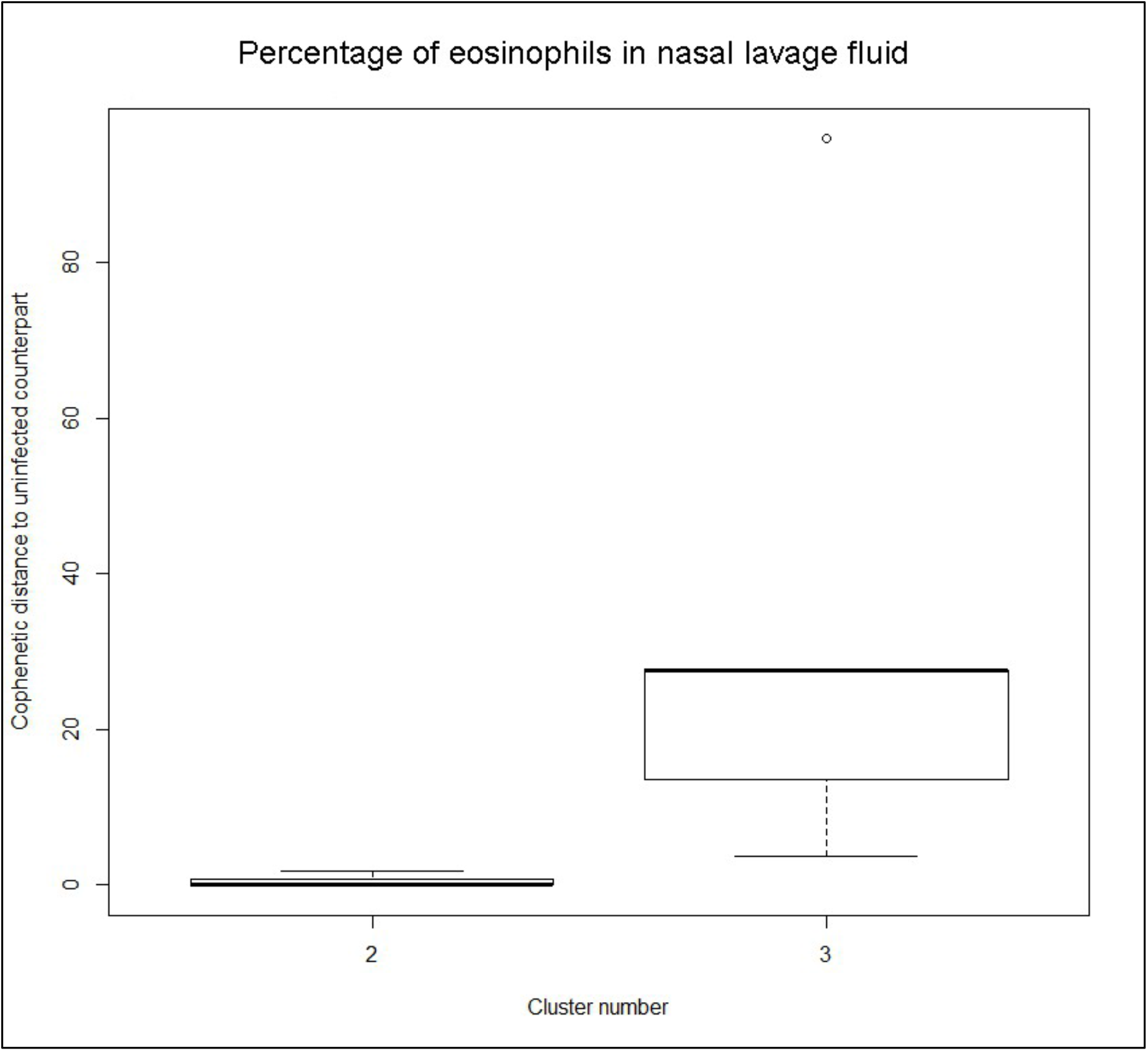
The boxplot to the left represents the distribution of cophenetic distances between time series corresponding to infected healthy participants and their uninfected counterparts. Only time series belonging to Cluster 2 in the clustering dendrogram obtained using the percentage of eosinophils in nasal lavage fluid (see Fig. 1 above) are contemplated here. The boxplot to the right represents the distribution of cophenetic distances between time series corresponding to infected asthmatic participants and their uninfected counterparts. Only time series belonging to Cluster 3 in the clustering dendrogram obtained using the percentage of eosinophils in nasal lavage fluid (see Fig. 1 above) are contemplated here. The two distributions are statistically significantly different (p-value= 8.96e-05, one-tailed Mann-Whitney-U-test, the cophenetic distances in Cluster 3 being, on average, higher than the ones in Cluster 2.

**Figure 3:**
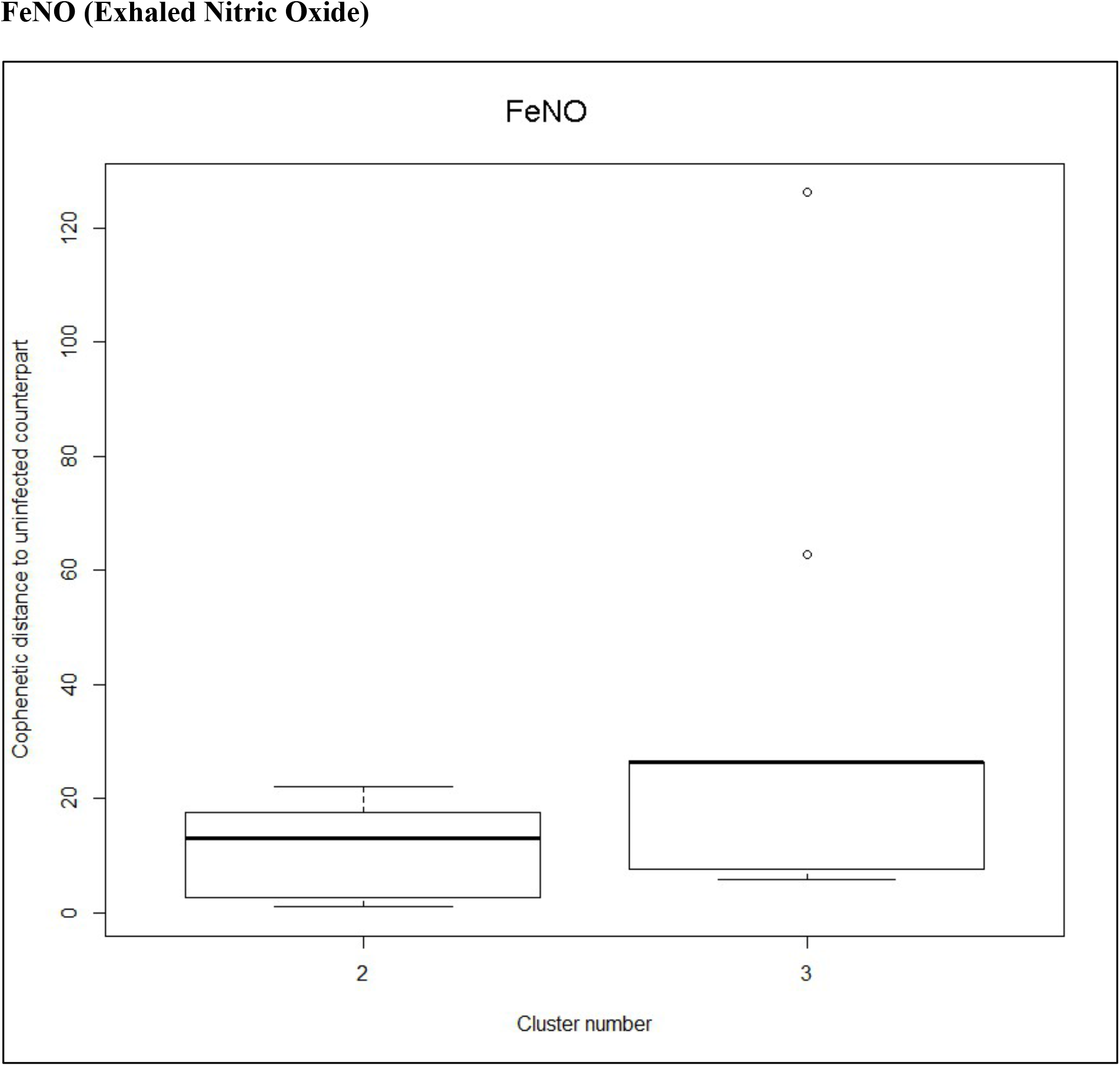
The boxplot to the left represents the distribution of cophenetic distances between time series corresponding to infected healthy participants and their uninfected counterparts. Only time series belonging to Cluster 2 in the clustering dendrogram obtained using FeNo data (see Fig. 1 in the Main Manuscript) are contemplated here. The boxplot to the right represents the distribution of cophenetic distances between time series corresponding to infected asthmatic participants and their uninfected counterparts. Only time series belonging to Cluster 3 in the clustering dendrogram obtained using FeNo data (see Fig. 1 in the Main Manuscript) are contemplated here. The two distributions are statistically significantly different (p-value=0.033, one-tailed Mann-Whitney-U-test, the cophenetic distances in Cluster 3 being, on average, higher than the ones in Cluster 2.

#### Cell density in nasal lavage fluid

We found one cluster of size 25 enriched in pre-challenge time series (p= 0.041), and one cluster of size 7 enriched in asthmatics (p=0.049). See Fig. 2

**Figure 4:**
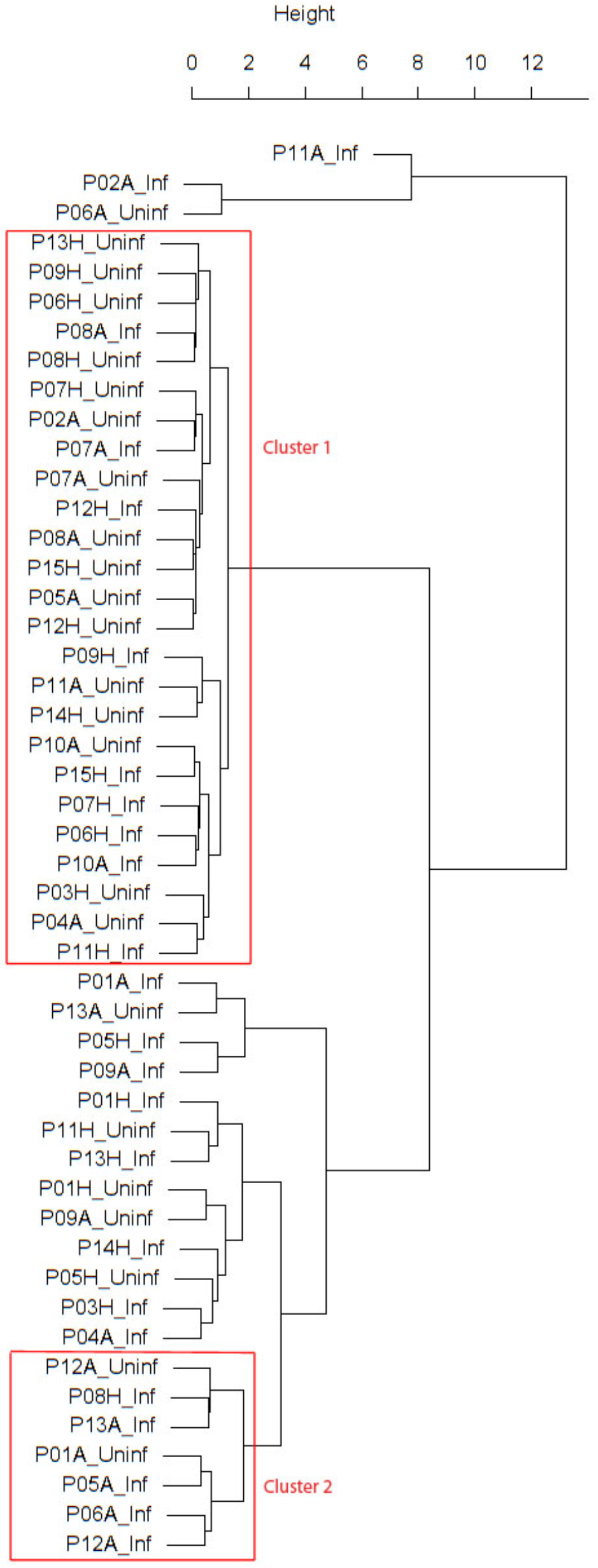
Dendrogram obtained from clustering the participants’ time series of cell density (millions per ml) in nasal lavage fluid using the EMD.

#### Percentage of neutrophils in nasal lavage fluid

We found one cluster of size 10 enriched in time series from healthy participants (p= 0.036). Moreover, no clusters were found showing a significant enrichment in or depletion of pre-challenge or post-challenge time series.

**Figure 5:**
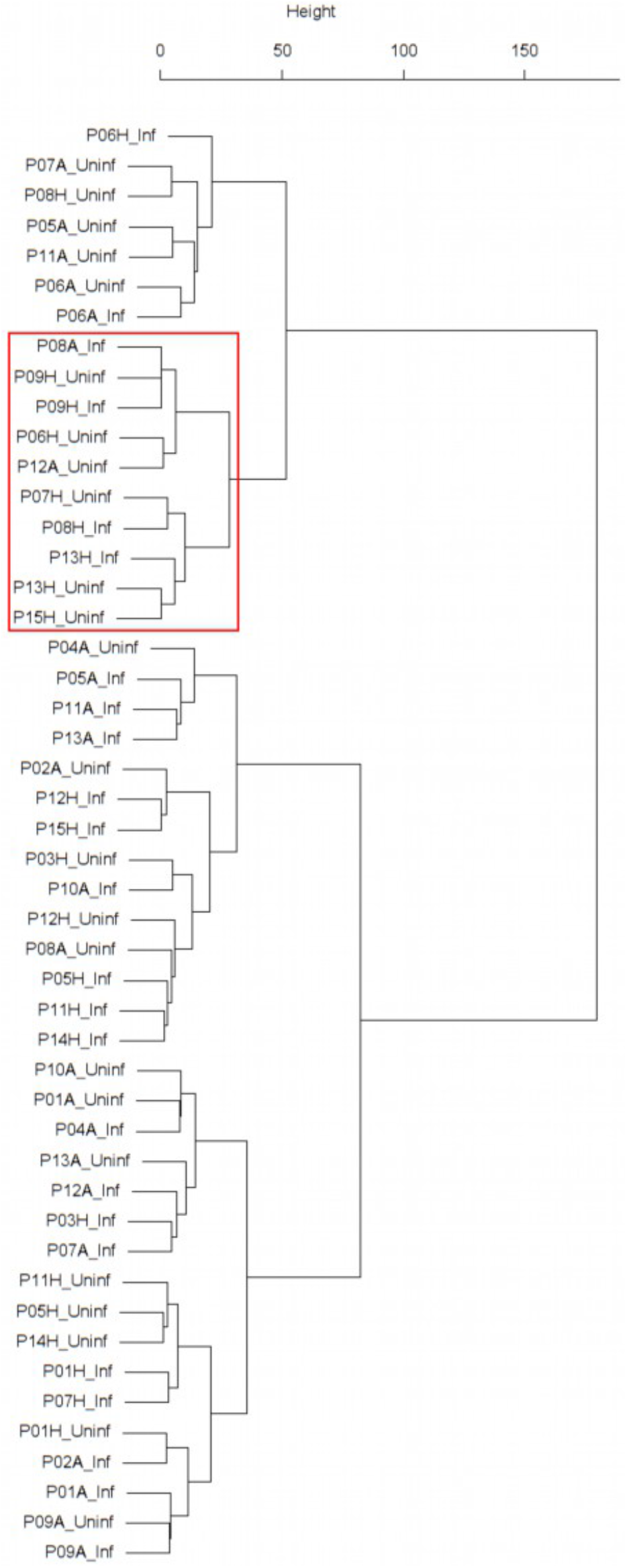
Dendrogram obtained from clustering the participants’ time series of the percentage of neutrophils in nasal lavage fluid using the EMD.

**Figure 6:**
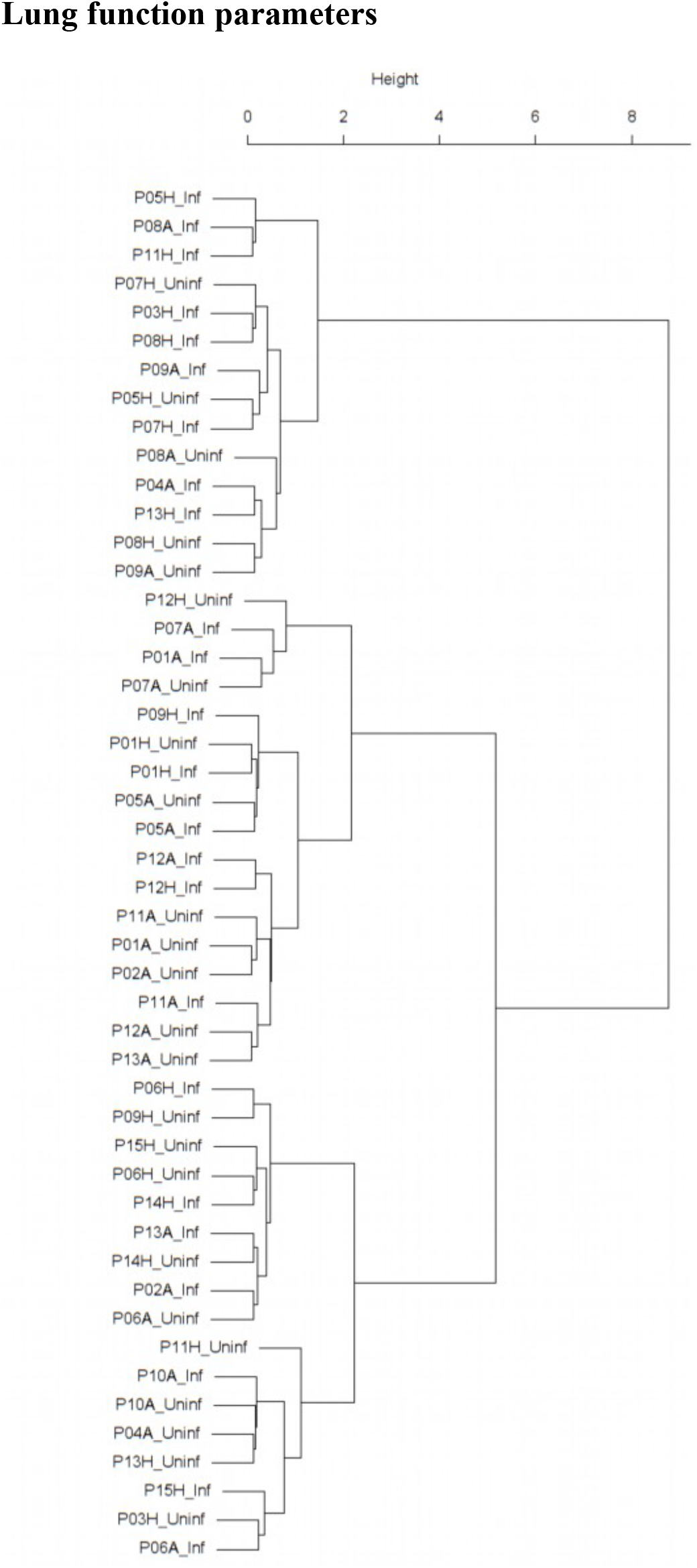
Dendrogram obtained from clustering the participants’ time series of the normalized ratio FEV1/FVC using the EMD.

**Figure 7:**
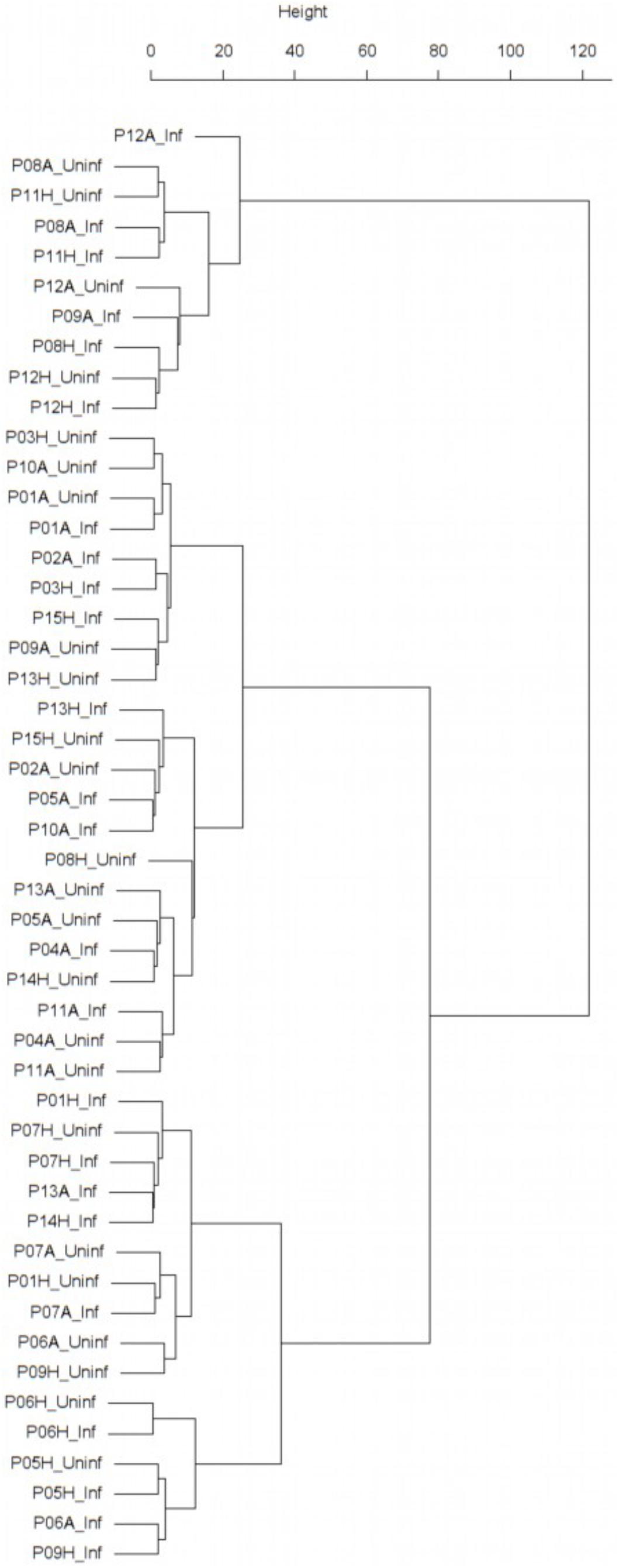
Dendrogram obtained from clustering the participants’ time series of PEF (% predicted) using the EMD.

**Figure 8:**
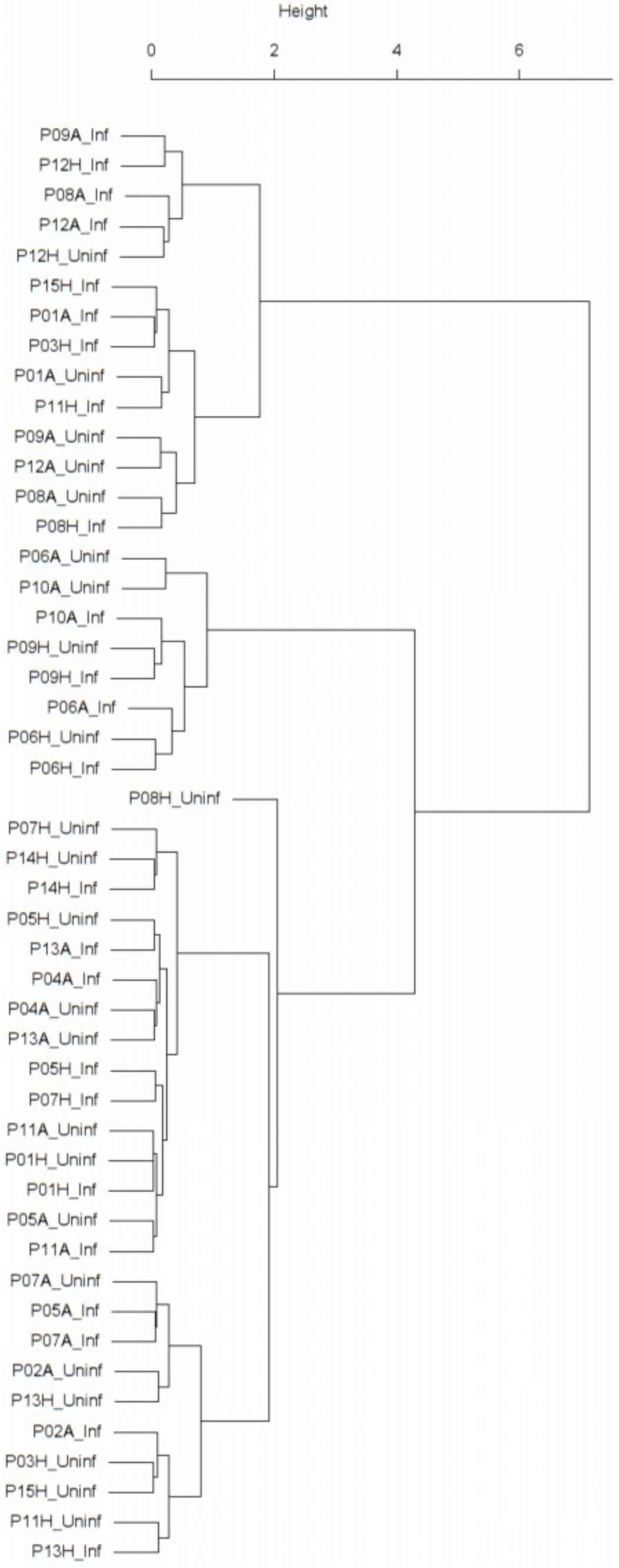
Dendrogram obtained from clustering the participants’ time series of normalized FEV1 using the EMD.

**Figure 9:**
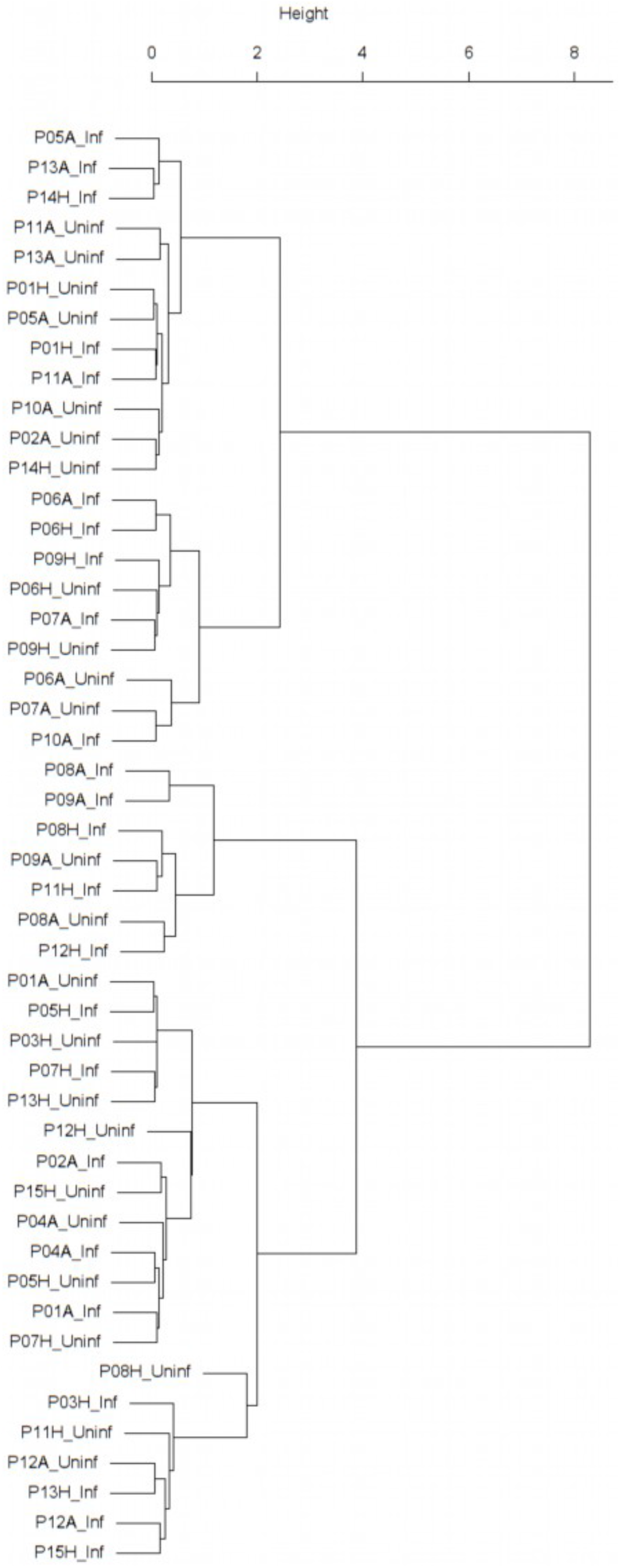
Dendrogram obtained from clustering the participants’ time series of normalized FVC using the EMD.

#### Sensitivity analysis of main clustering results

We investigated the sensitivity of the clustering of FeNO time series to changes in the data via nonparametric bootstrapping. However, given that the post-viral challenge time series are very short (eleven data points or fewer), resampling would be strongly affected by small sample size effects [1]. Thus, we only applied bootstrapping to the pre-challenge time series. Moreover, we resorted to soft bootstrapping [2] (see Methods below for more details) in order to increase the likelihood that least frequent values in the time series would be chosen during the resampling procedure. The results from 1000 soft bootstrapping iterations were as follows:

In 100% of the resulting soft bootstrap dendrograms, Cluster 1 (“outliers” cluster) was found. Moreover, two additional clusters, one, Cluster 2’, significantly enriched in time series stemming from heathy participants, and another, Cluster 3’, significantly enriched in time series stemming from asthmatic participants were found in 100% of the resulting soft bootstrap dendrograms. In other words, there was always a bootstrap counterpart to clusters 1,2, and 3 as found in the dendrogram obtained using the original, unperturbed data (see Figure 1 in the Main Manuscript).

In 51.4% of the soft bootstrap dendrograms, Cluster 3’ (the cluster enriched in time series stemming from asthmatic participants) contained a subcluster enriched in post-challenge time series. Whereas, only in 7.9% of the soft bootstrap dendrograms, Cluster 2’ (the cluster enriched in time series stemming from healthy participants) contained a subcluster enriched in post-challenge time series. (cf. Table 3 in the Main Manuscript).

Bootstrap distribution of mean cophenetic distances between the members of all pre- and post-pairs contained in Cluster 2’:

Min. 1st Qu. **Median** Mean 3rd Qu. Max.
6.358 11.017 **11.954** 13.527 14.947 26.654

Bootstrap distribution of mean cophenetic distances between the members of all pre- and post-pairs contained in Cluster 3’:

Min. 1st Qu. **Median** Mean 3rd Qu. Max.
7.036 21.581 **33.691** 29.321 35.610 49.080

In 80.6% of the soft bootstrap dendrograms, the mean cophenetic distances between the members of all pre- and post-pairs contained in Cluster 2’ was smaller than the mean cophenetic distances between the members of all pre- and post-pairs contained in Cluster 3’.

Bootstrap distribution of the p-values resulting from the one-tailed Mann-Whitney-U-test comparing the distribution of cophenetic distances between time series corresponding to infected healthy participants and their uninfected counterparts in Cluster 2’, to the distribution of cophenetic distances between time series corresponding to infected asthmatic participants and their uninfected counterparts in Cluster 3’ (cf. Figure 3 above):

Min. 1st Qu. **Median** Mean 3rd Qu. Max.
0.0005046 0.0163020 **0.0397435** 0.1563541 0.2208664 0.9537708

In 54.2% of the soft bootstrap dendrograms the resulting p-value was smaller or equal to 0.05. Bootstrap distribution of the percentage of neighboring pre- and post-pairs in Cluster 2’:

Min. 1st Qu. **Median** Mean 3rd Qu. Max.
0.00 14.29 **19.90** 19.13 23.53 42.86

Bootstrap distribution of the percentage of neighboring pre- and post-pairs in Cluster 3’:

Min. 1st Qu. **Median** Mean 3rd Qu. Max.
8.33 20.00 **25.00** 28.63 37.50 80.00

In 27.1% of the soft bootstrap dendrograms, the percentage of neighboring pre- and post-pairs in Cluster 2’was bigger than the percentage of neighboring pre- and post-pairs in Cluster 3’ (cf. Table 2 in the Main Manuscript).

Bootstrap distribution of the empirical p-values resulting from the permutation test used for establishing the statistical significance of the proportion of neighboring pre- and post-pairs found in Cluster 2’:

Min. 1st Qu. **Median** Mean 3rd Qu. Max.
0.00000 0.00596 **0.03780** 0.12118 0.15008 1.00000

In 52.7% of the soft bootstrap dendrograms, the resulting empirical p-value was smaller or equal to 0.05.

Bootstrap distribution of the empirical p-values resulting from the permutation test used for establishing the statistical significance of the proportion of neighboring pre- and post-pairs found in Cluster 3’:

Min. 1st Qu. **Median** Mean 3rd Qu. Max.
0.000000 0.004287 **0.014640** 0.080993 0.080477 0.450480

In 57.5% of the soft bootstrap dendrograms, the resulting empirical p-value was smaller or equal to 0.05.

Soft bootstrapping of the time series of the percentage of eosinophils in nasal lavage fluid yielded comparable results (data not shown).

## Materials and Methods

### 1. Participant cohort

24 study subjects were recruited among which 12 were healthy volunteers and the other 12 steroids naïve (or stopped using steroids 6 weeks prior to the study) mild to moderately persistent asthmatics.

The inclusion and exclusion criteria for participants to enter the study were as follows:

#### 1.1 Inclusion criteria

*Asthma patients* were selected using the following inclusion criteria:

- Age 18-50 years
- History of episodic chest tightness and wheezing
- Intermittent or mild to moderate persistent asthma according to the criteria by the Global Initiative for Asthma (Global Initiative of Asthma. www.ginasthma.org)
- Non-smoking or stopped smoking more than 12 months ago and 5 pack years or less
- Clinically stable, no exacerbations within last six weeks prior to study
- Steroid-naïve or those participants who are currently not on corticosteroids and have not taken any corticosteroids by any dosing-routes within 6 weeks prior to the study or only using on-demand reliever therapy
- Baseline pre-bronchodilator FEV_1_ ≥ 70% of predicted [3]
- Airway hyperresponsiveness, indicated by a positive acetyl-ß-methylcholine bromide (MeBr)challenge with PC_20_≤9.8 mg/ml [4]
- Positive skin prick test (SPT) to one or more of the 12 common aeroallergen extracts, defined as a wheal with an average diameter of ≥ 3 mm
- No other clinically significant abnormality on history and clinical examination
- Able to give written and dated informed consent prior to any study-specific procedures *Healthy subjects* were selected using the following inclusion criteria
- Age 18-50 years

- Non-smoking or stopped smoking more than 12 months ago and 5 pack years or less. Steroid-naïve, non-atopic participants who are currently not on any maintenance (subjects using oral contraceptives can be accepted)
- Baseline FEV_1_≥ 80% of predicted

- Negative acetyl-ß-methylcholine bromide (MeBr) challenge or PC_20_ ≥ 19.6 mg/ml
- Negative skin prick test (SPT) to one or more of the 12 common aeroallergen extracts
- Negative history of pulmonary and any other relevant disease
- Able to give written and dated informed consent prior to any study-specific procedure.

#### 1.2 Exclusion criteria

Potential subjects who meet any of the following criteria were excluded from participation in the study:

- Women who are pregnant, lactating or have a positive urine pregnancy test at baseline visit
- Participation in any clinical investigational drug treatment protocol within the preceding 5 half-lives (or 12 weeks, if the half-life is unknown) before the screening visit of this study.
- Concomitant disease or condition which could interfere with the conduct of the study, or for which the treatment might interfere with the conduct of the study, or which would, in the opinion of the investigator, pose an unacceptable risk to the patient

Furthermore, the following additional exclusion criteria were used in phase 2 of the study:

- RV16 titer > 1:8 in serum, measured at screening (visit 1) and also at the visit prior to the challenge
- History of clinically significant hypotensive episodes or symptoms of fainting, dizziness, or light-headedness
- History of an asthma exacerbation within the last 6 weeks prior to the study
- Has had any acute illness, including a common cold, within 4 weeks prior to visit 1
- Close contact with young children or with any immunosuppressed patients
- Has donated blood or has had a blood loss of more than 450 mL within 60 days prior to screening visit or plans to donate blood during the study
- Positive for any virus in nasal lavage at visit immediately prior to rhinovirus challenge.

#### 1.3 Sample size calculation

The study is explorative in nature and is based on some first estimates of fluctuating inflammatory biomarkers. A definite sample size for all biomarkers measured was not possible to be accurately determined.

A sample size of 12/12, based on previous studies with lesser data points, was calculated to identify biomarkers that have a strong, clinically relevant, sensitivity to the changes of disease stability over time [5, 6]. The sample size will probably have missed effects on biomarkers with weak effects, nevertheless, it has for all biomarkers screened provided evidence on long time variability, correlation and cross-correlation between biomarker properties, and susceptibility to strong viral external challenges. These three dimensions are a first estimate to understand the degree of variability and complexity of any disease process.

### 2. Study Design

The project represents a prospective observational, follow up study including patients with asthma and healthy controls with an experimental rhinovirus intervention in between.

#### 2.1 Screening Visit

First, the subjects were asked to sign an informed consent form after which they were examined for inclusion and exclusion criteria along with medical history and scoring of adverse events and concomitant medication. Spirometry, Methacholine challenge and Skin Prick Test along with physical examinations were performed meticulously.

#### 2.2 Run-in Phase

During this phase adverse events and medication were recorded along with physical examination. Exhaled nitric oxide (FeNO), nasal lavage, measurements were performed to acquaint the study subjects with the study procedures. Starting from the run-in, once-daily home-monitoring by Asthma Control Questionnaires, symptom scores and twice daily maneuvers by the pocket spirometer were executed.

#### 2.3 Baseline visit

The medication history (also adverse events and concomitant medication) was carefully recorded and thereby routine FeNO, and nasal lavage were measured. Pregnancy test was also performed.

#### 2.4 Study period

- **Phase 1:** The study participants were asked to visit the hospital clinic thrice weekly for the aforementioned measurements on Monday, Wednesday and Fridays, during the first 60 days.
- **Phase 2:** The participants were again followed up for the same measurements for the next 30 days with the same sampling frequency after being inoculated nasally with the common cold virus Rhinovirus 16, thereby mimicking the trigger for loss of asthma control or a mild exacerbation.

All participants in the study were screened for the presence of respiratory viruses just before the rhinovirus challenge, using Polymerase Chain Reaction confirmation on nasal lavage samples. This was performed in order to rule out a concomitant infection of the respiratory airways with viruses other than the one used in our study.

#### 2.5 End of study visit

All measurements were repeated at the end of each phase.

The schematic representation of the phases of the study is mentioned below in Fig. 10

**Figure 10:**
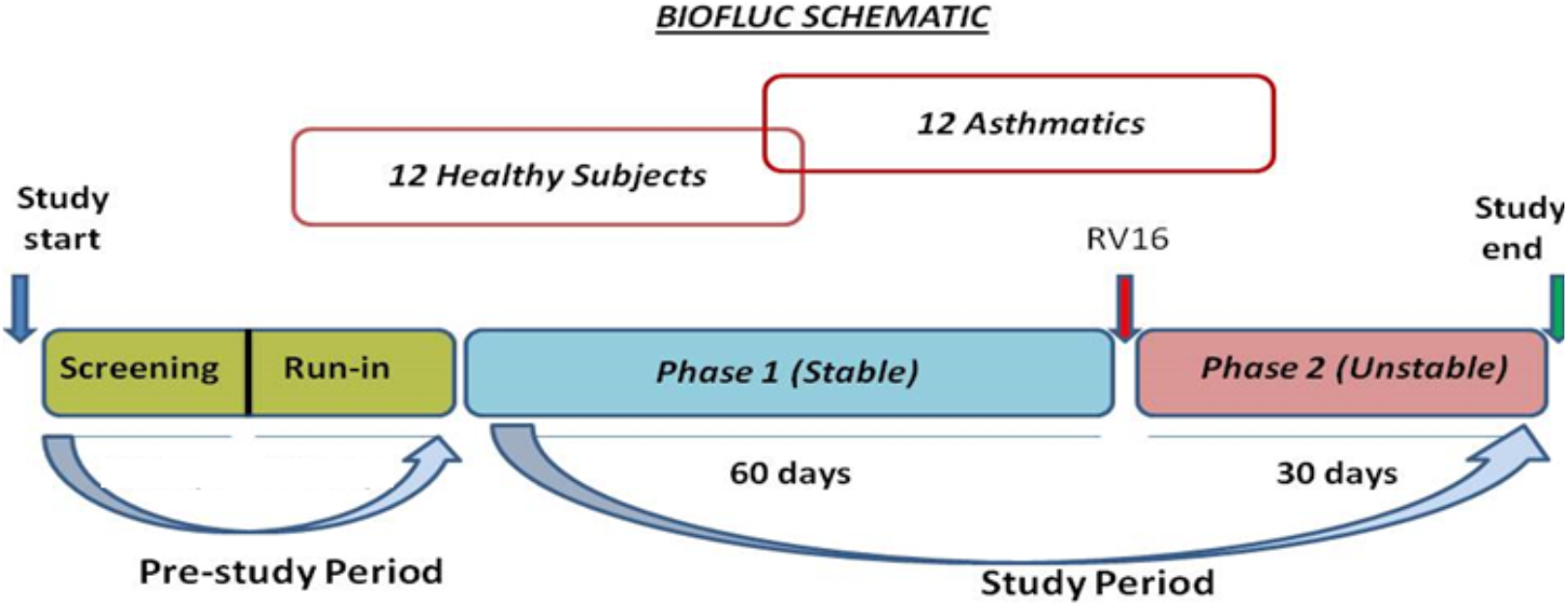
Schematic representation of the study design.

### 3. Measurement and collection of Biomarkers

#### 3.1 Study procedures

##### 3.1.1 Asthma Control Questionnaire

The Asthma Control Questionnaire (ACQ) was used in this study to assess the symptoms before and after Rhinovirus challenge along with lung function coupled to a hand-held spirometer device. Though widely used and well validated, it covers a 7-day time span and may not be optimal for a challenge protocol wherein symptoms change daily. Hence ACQ was used on a daily basis to record the daily changes in symptoms. Subjects complete the diary on rising in the morning and retiring at bedtime.

##### 3.1.2 Skin prick test

Allergy skin prick tests were performed based on the position paper by the European Academy of Allergology and Clinical Immunology (EAACI) [7]. The test was done by placing a drop of each of the 12 solutions containing common aeroallergens on the skin, followed by needle pricks by small SPT-lancets. Histamine was used as a positive control and saline solution as a negative control. The test result was considered positive if the skin develops a red, itchy area, called a ‘wheal’, with an average diameter of at least 3 mm, 15 minutes after the prick. The outline of all wheals was marked with a water-soluble marker and transferred to a test result from using adhesive tape, so as to facilitate measuring of the diameter of the wheal. All test result forms were archived. If the patient has been using anti-allergic medications, they were instructed to stop 3 days prior to the test in the clinic.

##### 3.1.3 Spirometry

Spirometry was performed using a daily calibrated spirometer according to European Respiratory Society (ERS) recommendations. Spirometry was done only once on the screening visit at the clinic. There was no use of bronchodilators (pre/post measurements) in the study. The subjects were asked not to use reliever medications at least 6 hours before the test starts in the clinic. An experienced lung function analyst from the AMC performed the lung function tests throughout the study to reach optimal performance and to enhance reproducibility. Spirometry (FEV1) results were printed out and documented. Home monitoring of morning and evening lung function were done by Peak Expiratory Flow Meter (PEF, Micro Diary, CareFusion).

Waking and bedtime FEV1 separately are of interest due to the exaggerated diurnal airflow variation seen in asthmatics. Subjects were encouraged to measure FEV1 at consistent times (upon waking between approximately 6am-9am and prior to bedtime between approximately 8pm-11pm, respectively – always prior to any bronchodilator therapy).

Forced expiratory flow 25–75% (FEF 25–75) was measured. Patients were systematically instructed to perform the home monitoring PEF measurements. The electronic PEF devices were tested and compared to the lab spirometry measurements.

###### Home monitoring

The morning and evening lung function measurements at home was done by Peak Expiratory Flow Meter (PEF, Micro Diary, CareFusion).

The following routine lung function indices were measured along with Asthma Control Questionnaire

FEV1: Forced Expiratory Volume in 1^st^ second
FVC: Forced Vital Capacity
FEV1/FVC: Ratio of the amount of air exhaled in 1^st^ second to the amount of air exhaled during a maximal expiration.
PEF: Peak Expiratory Flow

All the home recordings were monitored regularly (every 2 weeks) for uniformity and consistency of data. Stringent quality control of home lung function measurements was done at each visit.

##### 3.1.4 Exhaled nitric oxide

Measurements of fractionated exhaled nitric oxide (FENO) were performed (one measurement per subject) using the NIOX MINO^®^ (Aerocrine AB, Sweden). This analyser has an accuracy of ± 5 ppb or max 10%. The precision is < 3 ppb of a measured value < 30 ppb and ± 10 % of measured value > 30 ppb. Every individual subject was measured using a single measurement as standardised in the hospital.

Subjects were asked to perform a slow vital capacity manoeuvre with a standardised expiratory flow of 50 ml/sec for as long as possible. Positive mouth pressure was applied to close the velum and prevent contamination with NO from nose and paranasal sinuses. Expired gas was sampled continuously at the mouth piece and mean FENO values at the end expiratory plateau were calculated.

##### 3.1.5 Methacholine challenge

Methacholine is a cholinergic synthetic analogue of acetylcholine and acts directly on the airway smooth muscle resulting in bronchoconstriction. Measurement of airway responsiveness can be done using incremental inhaled doses of methacholine as bronchoprovocation test. This challenge test was performed using MeBr (acetyl-β-methylcholine bromide) according to the standardized tidal volume method that is operative in routine clinical diagnostics in the hospital [4].

##### 3.1.6 Nasal lavage

Nasal lavage was collected from the study participants once weekly before rhinovirus challenge and was up scaled to thrice weekly after the challenge at the clinic.

Eight ml of saline solution was introduced into the nasal cavity by a catheter and maintained for 10 min before recovery, followed by filtration and removal of mucous and cells by centrifugation. Standardized washings collected from the nose was used for biomarker analyses.

###### 3.1.6.1 Cell and slide preparation

A cell suspension of 200,000 cells/ml were prepared in PBS. 2 sets of Cytospin apparatus (Thermo Shandon Cytospin 4) were assembled and the cytospin filter was pre wet by spinning at 550 rpm for 1 minute with 50 μl PBS. 100 μl of the cell suspension was added to each slide and centrifuged at 450 rpm for 2 min. The quality of slides and the density of the cells were evaluated using a phase contrast microscopy. If the cell density was too high or low, more slides were prepared using an adjusted volume of the cell suspension. A target of 4 good slides with no overlapping or clumping of cells was set to improve the overall quality and consistency.

If the total cell count (for both slides together i.e. less than 7500 cells/slide) was less than or equal to 15000 cells, then the sample was centrifuged and re-suspended in 100 μl and was loaded onto a single block.

The prepared slides were stained as soon as possible with Diff Quick 1 stain (Medion Diagnostics), dried and mounted on Depex.

###### 3.1.6.2 Nasal cell cytology

A differential cell count on a maximum of 400 inflammatory cells were performed. Numbers of eosinophils, neutrophils, macrophages/monocytes, lymphocytes, columnar and squamous epithelial cells were recorded. A report was prepared with the overall counts, the quality of cytospin and additional aspects such as the presence of mucous, cell debris, inclusions, eosinophil granules etc.

##### 3.1.7 Blood venapunction

15 ml of venous blood was collected at each scheduled visit (once weekly) to determine whether circulating antibodies against RV16 are present, for hematology, and other immunological biomarker assays. Collection of blood was done in standard EDTA and non-EDTA tubes for specific purposes. 10 ml of it was immediately be followed by centrifugation to obtain plasma [2000g for 10 minutes (min) at room temperature, RT] aiding in removal of RBCs and WBCs. The supernatants were aliquoted for biomarker hunt.

#### 3.2 Rhinovirus challenge

In this study we exposed the volunteers to a mild dose (100 TCID 50) of RV16 using a validated approach which has been previously shown to be sufficient in inducing mild cold symptoms and decrease in lung function.

We used the GMP RV16 stock that has been tested previously by the medical ethical commission at the hospital in AUMC, and also by the U-BIOPRED showing efficacy in terms of cold symptoms and viral replication at 100 TCID50, which is part of the accompanying IMPD. This GMP RV16 stock has been prepared from a seed virus in extensively characterized human volunteers as described in METC 2010_310 and was expanded and aliquoted by Charles River Laboratories (USA) under GMP conditions. This preparation was considered safe for in vivo testing in human volunteers during a scientific advice meeting at BFarM (Bonn, Germany, April 30, 2013). This testing was carried out as per the approved protocol from the Amsterdam UMC medical ethical committee.

### 4. Statistical and computational Analysis

All computations were done using R [8] version 3.5.2 together with the following R-packages: car [9], reshape2 [10], openxlsx [11], lubridate [12], emdist [13], gplots [14], ape [15], ggdendro [16], cluster [17], factoextra [18], philentropy [19], dendextend [20], and plyr [21]. Statistical tests resulting in a p-value less or equal to 0.05 were regarded as significant.

#### 4.1 Assessment of differences: Pre- vs. post-viral-challenge

Consecutive measurements of a given biomarker prior to and after the viral challenge resulted in pre- and post-challenge time series for each cohort participant. Except for the calculation of transient changes, the time series were treated as empirical distributions, thus disregarding the chronological order of the measurements.

The empirical distribution of any biomarker before and after viral challenge was compared using Kolmogorov Smirnov test. A participant is considered a responder with respect to a given biomarker if the outcome of the Kolmogorov-Smirnov test results in a p-value <= 0.05. Differences in the variance between the pre- and post-challenge distributions were assessed using Levene’s test.

The empirical distributions of a given biomarker, prior to and after the viral challenge, were compared to each other using the Earth Mover’s Distance (EMD), see Fig. 2 in the Main Manuscript. The resulting pair-wise distances between distributions were then used for hierarchical clustering of pre- and post-challenge distributions. Our clustering approach makes use of the entire collection (distribution) of values measured before and after the challenge, respectively, and does not amalgamate the information into a single magnitude (e.g., the mean value). This method unveils subtler differences and similarities between the participants’ measurements that are less likely to be captured by conventional methods based on averages. Therefore, this part of our methodology is based on the distributional properties of each participant’s measurements and neglects the time dimension.

#### 4.2 Clustering approach

In our approach, for each biomarker, the measurements collected before and after the viral challenge were used to construct the individual empirical distribution of measurements for each study participant, before and after the challenge, respectively.

We performed agglomerative hierarchical clustering [22] of the aforementioned empirical distributions of biomarkers. Within the hierarchical clustering algorithm, the distances between the distributions were calculated using the Earth Mover’s Distance [23], and the agglomeration procedure was done according to Ward’s minimum variance method [22]. Intuitively speaking, the Earth Mover’s Distance contemplates the pair of distributions to be compared as piles of sand and measures the effort that it would take to shovel one distribution into the shape and position of the other (see below for a more detailed description of this method).

#### 4.3 Calculation of short-term/transient changes

For each participant individually, throughout the entire period of observation, the relative change of each biomarker taking place within time intervals of 10 days was calculated. This was done throughout the entire period of observation considering all possible time intervals consisting of 10 consecutive days, see Fig. 3 in the Main Manuscript. The rationale for this is as follows. Any changes taking place within a period of 10 days that contained the day of the viral challenge need to be interpreted and understood in the context of any changes taking place within a period of about 10 days during the entire pre-challenge phase of the study.

For each participant, the day of the challenge (i.e., the day of the inoculation) was labeled as “day 0”. All days between recruitment and the day of the challenge were labeled with negative integer numbers. All days between the day of the challenge and the participant’s final visit were labeled with positive integer numbers. Given that measurements were not conducted every day on a given participant (see study design above), for each biomarker considered in this study and for each participant separately, interpolation was used in order to have, for every biomarker one value for every day. Except for the total cell count in nasal lavage fluid, and the percentages of eosinophils and neutrophils in nasal lavage fluid, which were linearly interpolated, all biomarker time series were interpolated using cubic splines with natural boundary conditions (see, e.g., [24]). These interpolated values were only used for the assessment of the short-term/transient response.

The time interval of 10 days was decided based on prior knowledge where differences in signals from biomarkers (lung function etc.) usually take not less than a week to subside

As a sensitivity analysis, the same calculations were repeated using a 7-day time interval (data not shown). The outcomes were very similar to the ones obtained using the 10-day interval.

Consequently, a 10-day time period was considered optimal to calculate the short term/transient changes.

#### 4.4 Characterization of the dendrogram clusters

The clusters obtained using the clustering dendrogram were tested for enrichment in or depletion of healthy or asthmatic participants, and/or for enrichment in or depletion of pre- or post-challenge distributions.

The relative location of leaves in the clustering dendrogram was quantitatively evaluated using the cophenetic distance [25]. The cophenetic distance between two leaves of a dendrogram is defined as the height of the dendrogram at which the two largest branches that individually contain the two leaves merge into a single branch.

Two dendrogram leaves are called neighbors if their mutual cophenetic distance is equal to the minimum of all cophenetic distances from one of the leaves to all the other leaves in the dendrogram. If this condition is fulfilled for both leaves simultaneously, then the two leaves form a two-element cluster in the dendrogram. If the condition is only fulfilled for one of the leaves, the two are still considered neighbors, even if this is not always visually obvious from inspecting the dendrogram (see Fig. 11 below).

Under the null-hypothesis that the branching in the dendrogram is the result of a purely random process, the number of neighboring pre- and post-pairs to be expected just by chance within a given cluster can be estimated by simply permuting the labels of the leaves in the dendrogram and counting the number of neighboring pre- and post-pairs. This permutation test is used for calculating the empirical p-values displayed in Tables 1 and 3 in the Main Manuscript.

**Figure 11:**
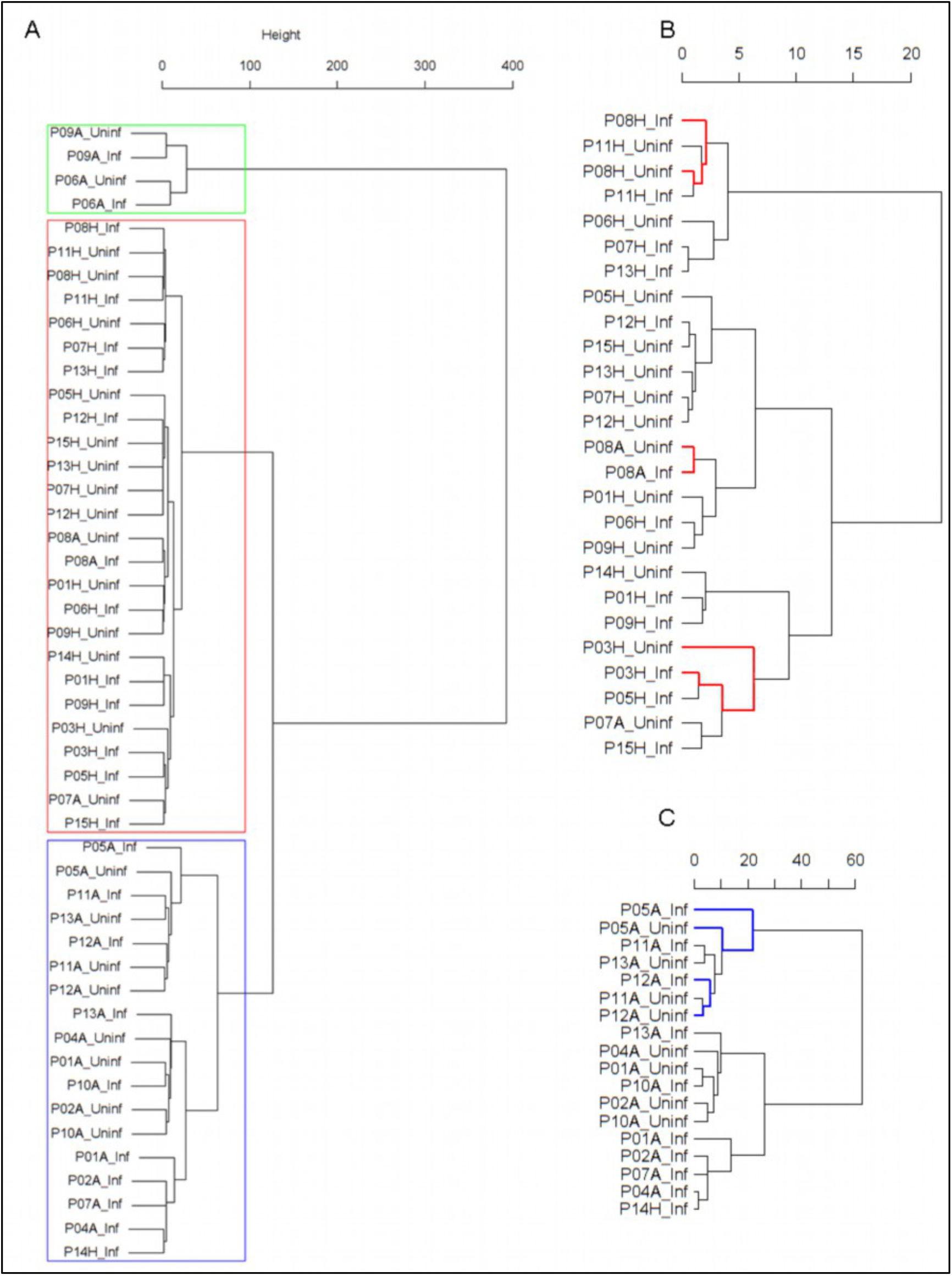
Panel A: Cluster dendrogram obtained via hierarchical clustering of the participants’ pre- and post-challenge time series of FeNO. The distance between any two-time series was calculated using the EMD. Rectangles mark the clusters identified. Panel B displays a more detailed view of the second cluster. According to the definition of neighboring leaves provided in the text above, the leaves P03H_Uninf and P03H_Inf are neighbors in this dendrogram (highlighted in red). The reason for this is that the cophenetic distance from leaf P03H_Uninf to leaf P03H_Inf is equal to the minimum of all distances from leaf P03H_Uninf to all other leaves in the dendrogram. This is not the case for the leaf P03H_Inf. However, the fact that this condition holds for at least one of the two leaves renders them neighboring. Leaves P08H_Inf and P08H_Uninf, and P11H_Inf and P11H_Uninf, are neighbors, respectively (the latter pair is not marked in color). P08A_Inf and P08A_Uninf are also neighbors; In this case, the minimum condition is fulfilled by both leaves for of this leaf-pair. This is why the two leaves form a two-element cluster in the dendrogram. Panel C displays a more detailed view of the third cluster. Analogous information about neighboring leaves in Cluster 3 is highlighted in blue. However, as opposed to Cluster 2 (depicted in Panel B), the amount of neighboring leaves in Cluster 3 is not statistically significant (permutation test, see Table 2 in the Main Manuscript).

#### 4.5 Soft Bootstrapping

As elucidated in [2], when resampling with replacement from a given time series in order to generate a bootstrap replicate of the same time series, the relative frequencies of the values in the original time series were adjusted in the sense of soft bootstrapping as follows:

1. The relative frequencies of the values in the original time series were sorted in ascending order: *p*_1_ ≤ *p*_2_ ≤ *p*_3_ ≤ ··· ≤ *p*_*m*_, where *m* is the number of different values in the time series.
2. The smallest (*p*_1_) and the strictly second-smallest (*p*_*q*_) relative frequencies were adjusted using a softness parameter (see [2]) *δ* = 0.008 according to the following formula:

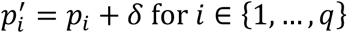
3. The remaining relative frequencies were adjusted according to the following formula:

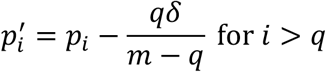

For time series with *m*<3 the corresponding relative frequencies were left unchanged during the soft bootstrapping iterations.

#### 4.6 The Earth Mover’s Distance

The Earth Mover’s Distance [23] is a method for quantifying the dissimilarity between two probability distributions. Intuitively speaking, the EMD contemplates the pair of distributions to be compared as piles of sand and measures the *minimal* effort that it would take to shovel one distribution into the shape and position of the other.

In practice, two distributions will be given by two representative samples, which can be written as lists of pairs {(*v*_*1*_, *w*_*1*_),…, (*v*_*n*_, *w*_*n*_)} and {(c_*1*_, *f*_*1*_),…, (c_*n*_, *f*_*m*_)}. Each pair (*v*_*i*_, *w*_*i*_) corresponds to a value *v*_*i*_ and its relative frequency *w*_*i*_ in the sample. If we translate the above described intuitive approach into numbers, the problem becomes a well-known transportation problem [26]: Suppose that *n* suppliers are located at positions *v*_*1*_,…, *v*_*n*_, respectively, and each one has a given amount of goods *w*_*i*_. Furthermore, they are required to supply *m* consumers, located at positions c_*1*_,…, c_*m*_, respectively, whereas each one has a given specific demand *f*_*i*_. For each supplier-consumer pair (*v*_*i*_, *w*_*i*_) and (c_*j*_, *f*_*j*_), the cost of transporting a single unit of goods is determined by the distance *d*(*v*_*i*_, *c*_*j*_) between their locations. The transportation problem is then to find a least expensive pattern of flow of goods from suppliers to consumers that would satisfy the consumers’ demand. Once the optimal pattern of the goods’ flow has been found, the total cost is the corresponding EMD.

Mathematically, this transportation problem can be formalized as a linear programming problem, for which efficient solution algorithms were developed in the late 1940s (see, e.g. [27]).

## Discussion

### Utility of biomarker time series analysis

Do longitudinal measurements provide deeper insights into complex disease physiology as compared to single measurements or average values? In order to answer this question, we tried to reproduce the results obtained using each participant’s entire collections of pre- and post-challenge biomarker measurements after collapsing them to the corresponding pre- and post-challenge individual average.

For example, in this cohort, FeNO time series have the ability to discriminate between healthy and asthmatics, and, within the group of asthmatics, between infected and uninfected (see Fig. 1 in the Main Manuscript). In order to investigate whether the average value would have a similar discriminative power, we calculated, for each participant, the average of their pre- and of their post-challenge series and used the absolute value of the difference between averages as a distance measure for clustering. The resulting dendrogram is depicted in Fig. 12 below. While the discriminative power between healthy and asthmatics is still given, the ability to distinguish between infected and uninfected within the group of asthmatics gets lost.

**Figure 12:**
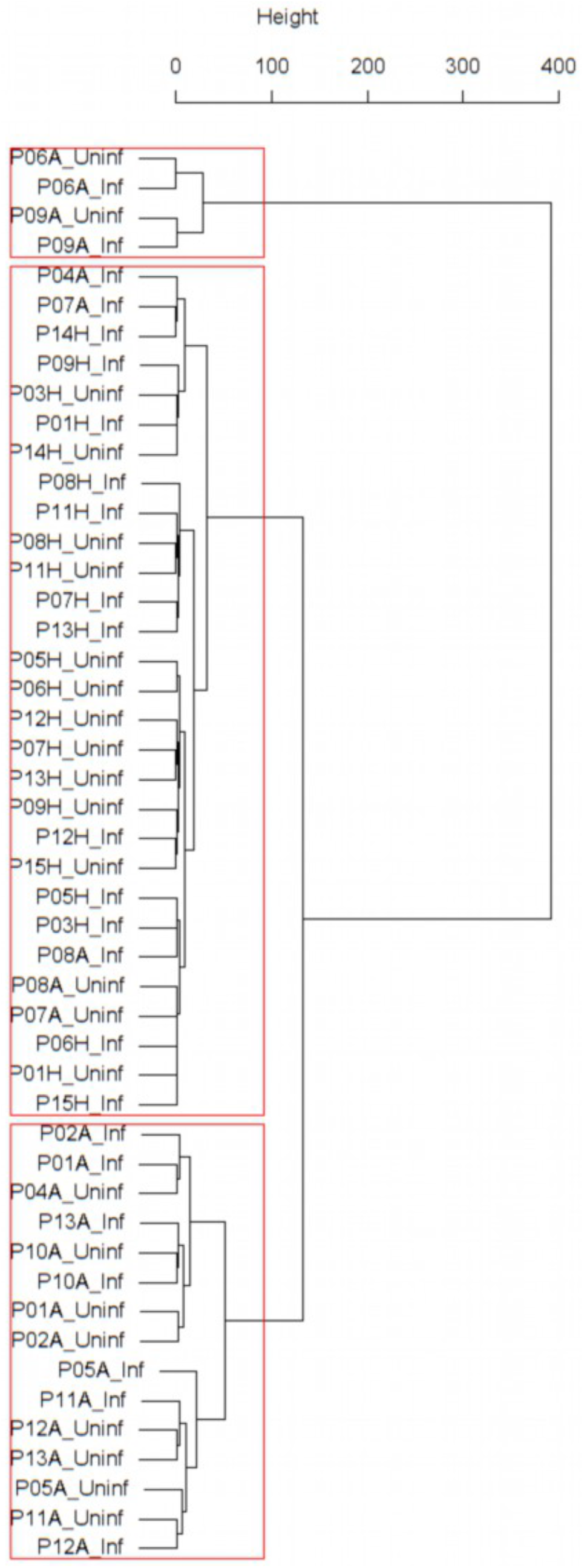
Dendrogram obtained from clustering the participants’ pre- and post-challenge *average* value of FeNO using the EMD.

## Summary Statistics of biomarkers in the study

### PEF (% predicted)

**Healthy group before viral challenge**

**Min. 1st Qu. Median Mean 3rd Qu. Max.**
**63.31 84.63 91.52 92.16 100.97 119.01**

**Healthy group after viral challenge**

**Min. 1st Qu. Median Mean 3rd Qu. Max.**
**63.42 78.42 91.92 90.44 102.39 118.84**

**Asthmatics group before viral challenge**

**Min. 1st Qu. Median Mean 3rd Qu. Max.**
**57.62 82.45 88.64 86.77 94.14 105.09**

**Asthmatics group after viral challenge**

**Min. 1st Qu. Median Mean 3rd Qu. Max.**
**47.67 78.49 88.44 85.01 95.21 109.98**

### Normalized FEV1

**Healthy group before viral challenge**

**Min. 1st Qu. Median Mean 3rd Qu. Max.**
**−2.7287 −1.5404 −0.9138 −1.0405 −0.7148 0.3588**

**Healthy group after viral challenge**

**Min. 1st Qu. Median Mean 3rd Qu. Max.**
**−3.0971 −1.9662 −1.3308 −1.3310 −0.8425 0.4305**

**Asthmatics group before viral challenge**

**Min. 1st Qu. Median Mean 3rd Qu. Max.**
**−2.4603 −2.1253 −1.0570 −1.3044 −0.8828 −0.1823**

**Asthmatics group after viral challenge**

**Min. 1st Qu. Median Mean 3rd Qu. Max.**
**−3.0350 −2.1484 −1.1480 −1.3369 −0.8418 0.5686**

### Normalized FVC

**Healthy group before viral challenge**

**Min. 1st Qu. Median Mean 3rd Qu. Max.**
**−2.19123 −1.87550 −1.66646 −1.34813 −0.83674 −0.02098**

**Healthy group after viral challenge**

**Min. 1st Qu. Median Mean 3rd Qu. Max.**
**−3.0841 −2.6730 −2.1174 −1.7966 −0.9592 0.2229**

**Asthmatics group before viral challenge**

**Min. 1st Qu. Median Mean 3rd Qu. Max.**
**−3.0455 −2.0029 −0.9102 −1.3416 −0.6539 −0.3614**

**Asthmatics group after viral challenge**

**Min. 1st Qu. Median Mean 3rd Qu. Max.**
**−3.726 −1.840 −1.319 −1.483 −0.668 0.194**

### Normalized FEV1/FVC

**Healthy group before viral challenge**

**Min. 1st Qu. Median Mean 3rd Qu. Max.**
**−1.4796 0.1851 0.6670 0.6940 1.3305 2.1131**

**Healthy group after viral challenge**

**Min. 1st Qu. Median Mean 3rd Qu. Max.**
**−0.6591 0.1088 1.3446 1.1083 1.9541 2.5255**

**Asthmatics group before viral challenge**

**Min. 1st Qu. Median Mean 3rd Qu. Max.**
**−1.2123 −0.5753 −0.3160 0.1677 1.2150 1.7450**

**Asthmatics group after viral challenge**

**Min. 1st Qu. Median Mean 3rd Qu. Max.**
**−1.5441 −0.4089 0.3237 0.4484 1.3195 2.5751**

### FeNO

**Healthy group before viral challenge**

**Min. 1st Qu. Median Mean 3rd Qu. Max.**
**9.435 12.779 13.872 14.608 15.594 22.227**

**Healthy group after viral challenge**

**Min. 1st Qu. Median Mean 3rd Qu. Max.**
**7.545 10.807 16.143 15.841 19.350 25.909**

**Asthmatics group before viral challenge**

**Min. 1st Qu. Median Mean 3rd Qu. Max.**
**16.57 35.67 46.21 61.97 55.68 181.30**

**Asthmatics group after viral challenge**

**Min. 1st Qu. Median Mean 3rd Qu. Max.**
**18.36 29.80 42.59 61.42 58.73 181.18**

### Cell density in 10^6^ per ml of nasal lavage fluid

**Healthy group before viral challenge**

**Min. 1st Qu. Median Mean 3rd Qu. Max.**
**0.1975 0.4308 0.5569 0.9327 1.2431 2.4600**

**Healthy group after viral challenge**

**Min. 1st Qu. Median Mean 3rd Qu. Max.**
**0.3436 0.6409 1.4536 1.6078 2.3982 3.7255**

**Asthmatics group before viral challenge**

**Min. 1st Qu. Median Mean 3rd Qu. Max.**
**0.0375 0.2490 0.8489 1.4183 1.8562 5.0444**

**Asthmatics group after viral challenge**

**Min. 1st Qu. Median Mean 3rd Qu. Max.**
**0.2873 0.9698 1.8445 2.7841 4.0850 10.8618**

### Percentage of Eosinophils in nasal lavage fluid

**Healthy group before viral challenge**

**Min. 1st Qu. Median Mean 3rd Qu. Max.**
**0.00000 0.00000 0.04375 0.95683 0.36979 9.03750**

**Healthy group after viral challenge**

**Min. 1st Qu. Median Mean 3rd Qu. Max.**
**0.0000 0.0000 0.0500 1.2977 0.2909 14.1273**

**Asthmatics group before viral challenge**

**Min. 1st Qu. Median Mean 3rd Qu. Max.**
**0.4091 7.2031 14.3950 16.9439 28.9778 35.9111**

**Asthmatics group after viral challenge**

**Min. 1st Qu. Median Mean 3rd Qu. Max.**
**0.000 7.218 10.227 13.186 13.935 59.736**

### Percentage of Neutrophils in nasal lavage fluid

**Healthy group before viral challenge**

**Min. 1st Qu. Median Mean 3rd Qu. Max.**
**24.90 47.36 58.90 62.45 86.44 99.64**

**Healthy group after viral challenge**

**Min. 1st Qu. Median Mean 3rd Qu. Max.**
**18.18 51.45 66.16 61.72 72.33 88.95**

**Asthmatics group before viral challenge**

**Min. 1st Qu. Median Mean 3rd Qu. Max.**
**29.30 36.18 60.47 57.23 79.79 92.72**

**Asthmatics group after viral challenge**

**Min. 1st Qu. Median Mean 3rd Qu. Max.**
**27.53 55.17 68.01 66.96 80.31 97.29**

For each biomarker, plots of the time series of relative changes within 10 days can be found in Supplementary Figures File 1.

Plots of the time series of each biomarker can be found in Supplementary Figures File 2.

## References

1. Chung KF. Clinical management of severe therapy-resistant asthma. Expert Review of Respiratory Medicine. 2017;11(5):395–402.

2. Frey U, Suki B. Complexity of chronic asthma and chronic obstructive pulmonary disease: implications for risk assessment, and disease progression and control. The Lancet. 2008;372(9643):1088–99.

3. Frey U, Maksym G, Suki B. Temporal complexity in clinical manifestations of lung disease. Journal of Applied Physiology. 2011;110(6):1723–31.

4. Macklem PT. Emergent phenomena and the secrets of life. Journal of Applied Physiology. 2008;104(6):1844–6.

5. Macklem PT, Seely A. Towards a definition of life. Perspect Biol Med. 2010;53(3):330–40.

6. Que C-L, Kenyon CM, Olivenstein R, Macklem PT, Maksym GN. Homeokinesis and short-term variability of human airway caliber. Journal of Applied Physiology. 2001;91(3):1131–41.

7. Yates FE. The 10th JAF Stevenson Memorial Lecture Outline of a physical theory of physiological systems. Canadian journal of physiology and pharmacology. 1982;60(3):217–48.

8. Garfinkel A. A mathematics for physiology. The American journal of physiology. 1983;245(4):R455–66.

9. Goldberger AL. Non-linear dynamics for clinicians: chaos theory, fractals, and complexity at the bedside. Lancet. 1996;347(9011):1312–4.

10. Goldberger AL, Amaral LA, Hausdorff JM, Ivanov P, Peng CK, Stanley HE. Fractal dynamics in physiology: alterations with disease and aging. Proceedings of the National Academy of Sciences of the United States of America. 2002;99 Suppl 1:2466–72.

11. Glass L. Synchronization and rhythmic processes in physiology. Nature. 2001;410:277.

12. Eke A, Herman P, Kocsis L, Kozak LR. Fractal characterization of complexity in temporal physiological signals. Physiol Meas. 2002;23(1):R1.

13. Amigó JM, Small M. Mathematical methods in medicine: neuroscience, cardiology and pathology. Philosophical transactions Series A, Mathematical, physical, and engineering sciences. 2017;375(2096):20170016.

14. Glass L, Kaplan D. Time series analysis of complex dynamics in physiology and medicine. Medical progress through technology. 1993;19:115–.

15. Glass L, Mackey MC. From clocks to chaos: the rhythms of life. Princeton, N.J.: Princeton University Press; 1988. xvii, 248 p. p.

16. Goldberger AL, Amaral LAN, Glass L, Hausdorff JM, Ivanov PC, Mark RG, et al. PhysioBank, PhysioToolkit, and PhysioNet: Components of a New Research Resource for Complex Physiologic Signals. Circulation. 2000;101(23):e215–e20.

17. Bélair J. Dynamical Disease: Mathematical Analysis of Human Illness: AIP Press; 1995.

18. Hall DM, Xu L, Drake VJ, Oberley LW, Oberley TD, Moseley PL, et al. Aging reduces adaptive capacity and stress protein expression in the liver after heat stress. Journal of Applied Physiology. 2000;89(2):749–59.

19. Orešic M, Vidal-Puig A. A systems biology approach to study metabolic syndrome. 2014.

20. Gan G, Ma C, Wu J. Data Clustering: Theory, Algorithms, and Applications: Society for Industrial and Applied Mathematics (SIAM, 3600 Market Street, Floor 6, Philadelphia, PA 19104); 2007.

21. Agresti A. A survey of exact inference for contingency tables. Statistical Science. 1992:131–53.

22. Fleiss JL, Levin B, Paik MC. Statistical Methods for Rates and Proportions: Wiley; 2013.

23. Rivals I, Personnaz L, Taing L, Potier M-C. Enrichment or depletion of a GO category within a class of genes: which test? Bioinformatics. 2007;23(4):401–7.

24. Johnston SL, Pattemore PK, Sanderson G, Smith S, Lampe F, Josephs L, et al. Community study of role of viral infections in exacerbations of asthma in 9-11 year old children. BMJ. 1995;310(6989):1225.

25. Nicholson KG, Kent J, Ireland DC. Respiratory viruses and exacerbations of asthma in adults. Br Med J. 1993;307(6910):982.

26. Su X, Ren Y, Li M, Zhao X, Kong L, Kang J. Prevalence of Comorbidities in Asthma and Nonasthma Patients: A Meta-analysis. Medicine (Baltimore). 2016;95(22):e3459–e.

27. Comberiati P, Katial RK, Covar RA. Bronchoprovocation Testing in Asthma: An Update. Immunology and Allergy Clinics of North America. 2018;38(4):545–71.

28. Bizzintino J, Lee WM, Laing IA, Vang F, Pappas T, Zhang G, et al. Association between human rhinovirus C and severity of acute asthma in children. Eur Respir J. 2011;37(5):1037–42.

29. Corne JM, Marshall C, Smith S, Schreiber J, Sanderson G, Holgate ST, et al. Frequency, severity, and duration of rhinovirus infections in asthmatic and non-asthmatic individuals: a longitudinal cohort study. The Lancet. 2002;359(9309):831–4.

30. Heymann PW, Carper HT, Murphy DD, Platts-Mills TAE, Patrie J, McLaughlin AP, et al. Viral infections in relation to age, atopy, and season of admission among children hospitalized for wheezing. Journal of Allergy and Clinical Immunology. 2004;114(2):239–47.

31. Johnston SL, Pattemore PK, Sanderson G, Smith S, Campbell MJ, Josephs LK, et al. The relationship between upper respiratory infections and hospital admissions for asthma: a time-trend analysis. American Journal of Respiratory and Critical Care Medicine. 1996;154(3):654–60.

32. Khetsuriani N, Lu X, Teague WG, Kazerouni N, Anderson LJ, Erdman DD. Novel human rhinoviruses and exacerbation of asthma in children. Emerging infectious diseases. 2008;14(11):1793–6.

33. Miller EK, Edwards KM, Weinberg GA, Iwane MK, Griffin MR, Hall CB, et al. A novel group of rhinoviruses is associated with asthma hospitalizations. Journal of Allergy and Clinical Immunology. 2009;123(1):98–104.e1.

34. Miller EK, Poehling KA, Edwards KM, Zhu Y, Griffin MR, Hartert TV, et al. Rhinovirus-Associated Hospitalizations in Young Children. The Journal of Infectious Diseases. 2007;195(6):773–81.

35. Murray CS, Poletti G, Kebadze T, Morris J, Woodcock A, Johnston SL, et al. Study of modifiable risk factors for asthma exacerbations: virus infection and allergen exposure increase the risk of asthma hospital admissions in children. Thorax. 2006;61(5):376.

36. Olenec JP, Kim WK, Lee W-M, Vang F, Pappas TE, Salazar LEP, et al. Weekly monitoring of children with asthma for infections and illness during common cold seasons. The Journal of allergy and clinical immunology. 2010;125(5):1001–6.e1.

37. Seemungal T, Harper-Owen R, Bhowmik A, Moric I, Sanderson G, Message S, et al. Respiratory Viruses, Symptoms, and Inflammatory Markers in Acute Exacerbations and Stable Chronic Obstructive Pulmonary Disease. American Journal of Respiratory and Critical Care Medicine. 2001;164(9):1618–23.

38. Smuts HE, Workman LJ, Zar HJ. Human rhinovirus infection in young African children with acute wheezing. BMC infectious diseases. 2011;11:65–.

39. Quint JK, Donaldson GC, Goldring JJ, Baghai-Ravary R, Hurst JR, Wedzicha JA. Serum IP-10 as a biomarker of human rhinovirus infection at exacerbation of COPD. Chest. 2010;137(4):812–22.

40. Wark PA, Bucchieri F, Johnston SL, Gibson PG, Hamilton L, Mimica J, et al. IFN-gamma-induced protein 10 is a novel biomarker of rhinovirus-induced asthma exacerbations. The Journal of allergy and clinical immunology. 2007;120(3):586–93.

41. Bardin PG, Fraenkel DJ, Sanderson G, van Schalkwyk EM, Holgate ST, Johnston SL. Peak expiratory flow changes during experimental rhinovirus infection. Eur Respir J. 2000;16(5):980–5.

42. de Kluijver J, Evertse CE, Sont JK, Schrumpf JA, van Zeijl-van der Ham CJ, Dick CR, et al. Are rhinovirus-induced airway responses in asthma aggravated by chronic allergen exposure? Am J Respir Crit Care Med. 2003;168(10):1174–80.

43. Rape M, Jentsch S. Taking a bite: proteasomal protein processing. Nature cell biology. 2002;4(5):E113–6.

44. Zhu J, Message SD, Qiu Y, Mallia P, Kebadze T, Contoli M, et al. Airway inflammation and illness severity in response to experimental rhinovirus infection in asthma. Chest. 2014;145(6):1219–29.

45. Miller MR, Hankinson J, Brusasco V, Burgos F, Casaburi R, Coates A, et al. Standardisation of spirometry. Eur Respir J. 2005;26(2):319–38.

46. Dweik RA, Boggs PB, Erzurum SC, Irvin CG, Leigh MW, Lundberg JO, et al. An official ATS clinical practice guideline: interpretation of exhaled nitric oxide levels (FENO) for clinical applications. Am J Respir Crit Care Med. 2011;184(5):602–15.

47. DeMore JP, Weisshaar EH, Vrtis RF, Swenson CA, Evans MD, Morin A, et al. Similar colds in subjects with allergic asthma and nonatopic subjects after inoculation with rhinovirus-16. The Journal of allergy and clinical immunology. 2009;124(2):245–52, 52 e1-3.

48. Gern JE, Calhoun W, Swenson C, Shen G, Busse WW. Rhinovirus infection preferentially increases lower airway responsiveness in allergic subjects. Am J Respir Crit Care Med. 1997;155(6):1872–6.

49. Grunberg K, Sharon RF, Sont JK, In ‘t Veen JC, Van Schadewijk WA, De Klerk EP, et al. Rhinovirusinduced airway inflammation in asthma: effect of treatment with inhaled corticosteroids before and during experimental infection. Am J Respir Crit Care Med. 2001;164(10 Pt 1):1816–22.

50. Grunberg K, Timmers MC, de Klerk EP, Dick EC, Sterk PJ. Experimental rhinovirus 16 infection causes variable airway obstruction in subjects with atopic asthma. Am J Respir Crit Care Med. 1999;160(4):1375–80.

51. Mallia P, Message SD, Kebadze T, Parker HL, Kon OM, Johnston SL. An experimental model of rhinovirus induced chronic obstructive pulmonary disease exacerbations: a pilot study. Respir Res. 2006;7:116.

52. Traves SL, Proud D. Viral-associated exacerbations of asthma and COPD. Curr Opin Pharmacol. 2007;7(3):252–8.

53. Ravi A, Koster J, Dijkhuis A, Bal SM, Sabogal Piñeros YS, Bonta PI, et al. Interferon-induced epithelial response to rhinovirus-16 in asthma relates to inflammation and FEV1. Journal of Allergy and Clinical Immunology. 2018.

54. Levene H. Robust tests for equality of variances. Contributions to Probability and Statistics Essays in Honor of Harold Hotelling. 1961:279–92.

55. Benjamini Y, Hochberg Y. Controlling the False Discovery Rate: A Practical and Powerful Approach to Multiple Testing. Journal of the Royal Statistical Society Series B (Methodological). 1995;57(1):289–300.

56. Rubner Y, Tomasi C, Guibas LJ, editors. A metric for distributions with applications to image databases. Computer Vision, 1998 Sixth International Conference on; 1998: IEEE.

57. Denlinger LC, Sorkness RL, Lee W-M, Evans MD, Wolff MJ, Mathur SK, et al. Lower Airway Rhinovirus Burden and the Seasonal Risk of Asthma Exacerbation. American Journal of Respiratory and Critical Care Medicine. 2011;184(9):1007–14.

58. Sokal RR, Rohlf FJ. The Comparison of Dendrograms by Objective Methods. Taxon. 1962;11(2):33–40.

## References

1. Isaksson A, Wallman M, Göransson H, Gustafsson MG. Cross-validation and bootstrapping are unreliable in small sample classification. Pattern Recognition Letters. 2008;29(14):1960–5. doi: https://doi.org/10.1016/j.patrec.2008.06.018.

2. Mucha H-J, Bartel H-G, editors. Soft Bootstrapping in Cluster Analysis and Its Comparison with Other Resampling Methods. Data Analysis, Machine Learning and Knowledge Discovery; 2014 2014//; Cham: Springer International Publishing.

3. Quanjer PH, Tammeling GJ, Cotes JE, Pedersen OF, Peslin R, Yernault JC. Lung volumes and forced ventilatory flows. Report Working Party Standardization of Lung Function Tests, European Community for Steel and Coal. Official Statement of the European Respiratory Society. Eur Respir J Suppl. 1993;16:5–40. Epub 1993/03/01. PubMed PMID: 8499054.

4. Sterk PJ, Fabbri LM, Quanjer PH, Cockcroft DW, O’Byrne PM, Anderson SD, et al. Standardized challenge testing with pharmacological, physical and sensitizing stimuli in adults. Eur Respir J. 1993;6 Suppl 16:53–83. Epub 1993/03/01. doi: 10.1183/09041950.053s1693. PubMed PMID: 24576917.

5. Turner C, Spanel P, Smith D. A longitudinal study of ethanol and acetaldehyde in the exhaled breath of healthy volunteers using selected-ion flow-tube mass spectrometry. Rapid Commun Mass Spectrom. 2006;20(1):61–8. Epub 2005/11/29. doi: 10.1002/rcm.2275. PubMed PMID: 16312013.

6. Turner C, Spanel P, Smith D. A longitudinal study of breath isoprene in healthy volunteers using selected ion flow tube mass spectrometry (SIFT-MS). Physiol Meas. 2006;27(1):13–22. Epub 2005/12/21. doi: 10.1088/0967-3334/27/1/002. PubMed PMID: 16365507.

7. Position paper: Allergen standardization and skin tests. The European Academy of Allergology and Clinical Immunology. Allergy. 1993;48(14 Suppl):48–82. Epub 1993/01/01. PubMed PMID: 8342740.

8. R Core Team. R: A Language and Environment for Statistical Computing Vienna, Austria: R Foundation for Statistical Computing; 2018. Available from: https://www.R-project.org.

9. Fox J, Weisberg S. An {R} Companion to Applied Regression. Second ed: Thousand Oaks; 2011.

10. Wickham H. Reshaping Data with the reshape Package. Journal of Statistical Software; Vol 1, Issue 12 (2007). 2007.

11. Walker A. openxlsx: Read, Write and Edit XLSX Files. R package version 4.1.0. https://CRAN.R-project.org/package=openxlsx. 2018.

12. Grolemund G, Wickham H. Dates and Times Made Easy with lubridate. Journal of Statistical Software; Vol 1, Issue 3 (2011). 2011.

13. Urbanek S, Rubner Y. emdist: Earth Mover’s Distance. 2012.

14. Warnes GR, Bolker B, Bonebakker L, Gentleman R, Liaw WHA, Lumley T, et al. gplots: Various R programming tools for plotting data. 2014.

15. Paradis E, Claude J, Strimmer K. APE: analyses of phylogenetics and evolution in R language. Bioinformatics. 2004;20:289–90.

16. ggdendro: Tools for extracting dendrogram and tree diagram plot data for use with ggplot. [Internet]. 2013. Available from: http://CRAN.R-project.org/package=ggdendro.

17. Maechler M, Rousseeuw P, Struyf A, Hubert M, Hornik K. cluster: Cluster Analysis Basics and Extensions. R package version 2.0.7-1. 2018.

18. Kassambara A, Mundt F. factoextra: Extract and Visualize the Results of Multivariate Data Analyses. R package version 1.0.5. https://CRAN.R-project.org/package=factoextra. 2017.

19. Drost H-G. philentropy: Similarity and Distance Quantification Between Probability Functions. R package version 0.2.0. https://CRAN.R-project.org/package=philentropy. 2018.

20. Galili T. dendextend: an R package for visualizing, adjusting and comparing trees of hierarchical clustering. Bioinformatics (Oxford, England). 2015;31(22):3718–20. Epub 2015/07/23. doi: 10.1093/bioinformatics/btv428. PubMed PMID: 26209431.

21. Wickham H. The Split-Apply-Combine Strategy for Data Analysis. Journal of Statistical Software; Vol 1, Issue 1 (2011). 2011. doi: 10.18637/jss.v040.i01.

22. Gan G, Ma C, Wu J. Data Clustering: Theory, Algorithms, and Applications: Society for Industrial and Applied Mathematics (SIAM, 3600 Market Street, Floor 6, Philadelphia, PA 19104); 2007.

23. Rubner Y, Tomasi C, Guibas LJ, editors. A metric for distributions with applications to image databases. Computer Vision, 1998 Sixth International Conference on; 1998: IEEE.

24. Bartels RH, Beatty JC, Barsky BA. An introduction to splines for use in computer graphics and geometric modeling: Morgan Kaufmann; 1995.

25. Sokal RR, Rohlf FJ. The Comparison of Dendrograms by Objective Methods. Taxon. 1962;11(2):33–40. doi: 10.2307/1217208.

26. Hitchcock FL. The Distribution of a Product from Several Sources to Numerous Localities. Journal of Mathematics and Physics. 1941;20(1-4):224–30. doi: 10.1002/sapm1941201224.

27. Hillier FS, Lieberman GJ. Introduction to operations research. 9th ed. New York: McGraw-Hill Higher Education; 2010. xxiv, 1047 p. p.

